# Effective recognition of double-stranded RNA does not require activation of cellular inflammation

**DOI:** 10.1101/2024.05.24.595663

**Authors:** Karolina Drazkowska, Julia Cieslicka, Anna Pastucha, Lukasz Markiewicz, Krzysztof Goryca, Tomasz Kowalczyk, Dominik Cysewski, Andreas Bausch, Pawel J. Sikorski

## Abstract

Excess double-stranded RNA (dsRNA) is present in the cytoplasm of human cells, usually following viral infections. Recognition of dsRNAs activates innate immune pathways, leading to cellular inflammation and inhibition of cell growth. Here, we show that an effective dsRNA response may occur without the onset of inflammation. Interestingly, pro-inflammatory (RLR-dependent pathway) and cell growth inhibitory mechanisms (OAS/RNase L- and PKR-dependent pathways) can act independently. We found that the 5′ ends of dsRNA direct the onset of cellular inflammation, whereas RNA duplex activates the OAS/RNase L and PKR pathways. Unexpectedly, three of the most common human RNA epitranscriptomic markers – *i.e. N*6-methyladenosine, 5-methylcytosine, and pseudouridine – did not affect the immunogenicity of dsRNA; however, the presence of *N*6-methyladenosine inhibited the OAS/RNase L pathway. Our observations demonstrate how precisely innate immunity is fine-tuned in cells to take appropriate countermeasures when a specific threat arises.

## Introduction

Double-stranded RNA (dsRNA) in human cells originates from endogenous (self-derived) and exogenous sources.^1,2^ Healthy human cells growing under optimal conditions have low levels of dsRNAs derived from, for example, the RNA interference pathway or are related to the presence of long non-coding RNA and genomic transposable elements. However, in some pathological conditions, dysregulation of the RNA machinery leads to the accumulation of excessive amounts of dsRNA.^1,3–8^ Viral infection is the primary source of dsRNA in human cells. In such situations, the dsRNA present in the host cell cytoplasm may be either an intermediate product of viral replication or the viral genome itself; some viral genomes consist of dsRNA.^2,9^ Therefore, human cells have evolved intricate mechanisms to recognize and respond to the presence of dsRNA as a potential threat of viral infection. This recognition is critical for triggering the human innate immune response, which is the first line of defense against viral infections.

Upon recognition of dsRNA, the host cell initiates an antiviral response aimed at limiting viral replication as much as possible and alerting neighboring uninfected cells of viral threats. In the cytoplasm of the host cell, dsRNAs are recognized by retinoic acid-inducible gene I (RIG-I)-like receptors (RLRs), which activate inflammatory signaling pathways through the mitochondrial antiviral signaling (MAVS) adaptor protein, leading to the production of interferons and cytokines and the activation of interferon-stimulated genes (ISGs).^10–13^ In addition, host cells attempt to limit viral replication by slowing down their own metabolism and sacrificing their well-being. Two main mechanisms inhibit host cell growth upon viral infection: dsRNA-activated protein kinase (PKR) and oligoadenylate synthetase (OAS)/RNase L-dependent pathways. Activation of the host PKR leads to a global shutdown of protein biosynthesis, resulting in reduced viral and host protein production.^14–16^ In contrast, viral and host single-stranded transcripts are degraded upon activation of the OAS/RNase L pathway.^17–19^ Host sensors differ in their mechanism of dsRNA recognition, recognizing specific dsRNA features or simply detecting the presence of a duplex of ribonucleic acids. RIG-I recognizes dsRNA based on the dsRNA 5’ end structure and has the highest affinity for triphosphorylated dsRNA (ppp-dsRNA).^11,13,20–22^ It was thought that the presence of the cap structure was sufficient to significantly decrease the affinity of RIG-I for dsRNA, but dsRNA with cap0 had similar affinity for dsRNA as the triphosphorylated counterpart, and only 2’-*O*-methylation of the cap structure abolishes the RIG-I binding to dsRNA.^20^ Another member of the RLR family, melanoma differentiation-associated protein 5 (MDA5), recognizes long duplex structures within transcripts rather than 5’ end modifications present on dsRNAs.^23,24^ Other reports showed that 5’ end modifications of dsRNA may play a role in its recognition by MDA5.^12,25^ However, the exact mechanism by which MDA5 senses dsRNA is not yet fully understood. In the case of OAS1/2/3 and PKR, the presence of the RNA duplex seems to be the only requirement to classify the transcripts as a potential threat to the host cell.^15,26,27^

Viruses, in the course of evolution, have acquired mechanisms that allow them to shield their nucleic acids from recognition by host sensors.^2,9^ To avoid recognition by receptors that sense the 5’ end of dsRNA, viral RNA (vRNA) can be capped like endogenous human transcripts.^28^ The cap structure not only prevents RIG-I from binding to the vRNA, but also allows viruses to benefit from cap-dependent translation. In addition, vRNAs could undergo post-transcriptional modifications, similar to those observed in host transcripts, to mimic endogenous RNA molecules as closely as possible and further evade immune recognition.^29^ Among the modifications that have been identified in viral RNA are those that are most abundant in the human transcriptome, namely *N*6-methyladenosine (m^6^A), pseudouridine (Ψ), and 5-methylcytosine (m^5^C).

Although viral strategies to evade immune recognition have been known for many years and the innate immune pathways leading to the activation of antiviral responses and inhibition of host cell growth have been thoroughly characterized, we still do not know how each feature of dsRNA affects host cell defense mechanisms. Does the mere presence of the RNA duplex, regardless of how it is modified, activate all the innate immune pathways of the host cell, or is each specific feature of the dsRNA responsible for activating a particular defense mechanism? Is it possible that, under certain conditions, these pathways are so independent of each other that dsRNA can activate only some of them? Does dsRNA recognition always lead to the activation of all possible defense mechanisms? Studies on the response of human cells to viral infection have always led to the conclusion that the presence of viral RNA activates all possible defense mechanisms to fight the infection.^2,9^ A similar answer is provided by studies using a synthetic polyinosinic-polycytidylic acid compound, better known as poly(I:C), which is widely used to mimic viral infection.^27,30–35^ More detailed information about the function of individual host sensors comes from *in vitro* studies; however, based on these results, it is difficult to understand how the innate defense system as a whole functions and how it is regulated.

Herein, we employed a novel approach and used an *in vitro* prepared dsRNA carrying various modifications both at the 5’ end and in its body, or combinations thereof, to truly characterize host cell-dsRNA interplay. The use of properly prepared and modified dsRNAs allowed us to understand how this class of transcripts activated the innate immune pathway. We revealed that the activation of the immune pathway leading to the inhibition of cell growth can be achieved without the onset of cell inflammation. We showed that modifications at the 5’ end of dsRNA only affect the MAVS-dependent antiviral and inflammatory signaling pathway. The activation of the OAS/RNase L and PKR pathways, leading to cell growth inhibition, is exclusively dependent on the detection of ribonucleic acid duplexes, and their activation does not lead to interferon production. In addition, we outlined the extent to which epitranscriptomic markers affect innate immune pathways and found that m^6^A modification in the body of transcripts can protect dsRNAs from recognition by OAS proteins, whereas none of the tested modifications affected dsRNA recognition by PKR.

## Results

### Cap structure is the key feature of transcripts responsible for dsRNA immunogenicity

The most widely used synthetic molecule to mimic viral infection and trigger the innate immune system is the poly(I:C) molecule.^27,30–35^ This synthetic polymer not only activates RLR pathways but also stimulates OAS/RNase L and PKR-dependent defense mechanisms. Recently, we and others have shown that dsRNA generated by *in vitro* transcription using T7 RNA polymerase can be used to produce molecules with immunostimulatory potential similar to poly(I:C).^35–37^ Importantly, these dsRNA molecules have a triphosphate group at their 5’ ends, and it is widely accepted that the immunogenicity of dsRNA is due to the presence of this chemical group.^11,12,21,23,36^ Therefore the question arises: how the presence of a cap at the 5’ ends of dsRNA modulates its immunogenic potential? In addition, mammalian transcripts have different versions of the cap structure that differ in the amount of 2’-*O*-methyl group attached to the 5’ end of RNA. On this basis we can distinguish cap0, cap1, and cap2 (cap0 is unmethylated, cap1 has one 2’-*O*-methylation at the first transcribed nucleotide, and cap2 has two methyl groups at the first and second transcribed nucleotide). Recent studies on mammalian messenger RNAs (mRNAs) revealed that 2’-*O*-methylation(s) within cap label transcripts as self-ones, thus cell treats triphosphorylated single-stranded RNAs (ssRNAs) or these ones with cap0 as a potential threat. Interestingly, RNA viruses in the course of evolution have acquired the ability not only to add a cap structure to their RNAs, but also to generate their transcripts with a 2’-*O*-methylated version of the cap, *i.e.* transcripts with cap1. Does this mean that dsRNAs formed during the replication of such viruses avoid immune recognition inside human cells? Moreover, if 2’-*O*-methylation(s) reduce(s) the immunogenicity of the dsRNA, does this mean that only dsRNA with cap0 is immunogenic? A previous report showed that short blunt-ended dsRNA with cap0 bound to RIG-I with almost the same affinity as ppp-dsRNA,^20^ therefore, does this mean that dsRNA with cap0 triggers the innate immune response to the same extent as ppp-dsRNA? To answer these questions, we prepared a set of dsRNAs that differ in their 5’ end modifications. This was made possible by taking advantage of *in vitro* transcription reactions. By using appropriate cap analogs for RNA synthesis, we were able to generate ssRNAs with the cap structure of interest at their 5’ ends.^36,38^ To obtain dsRNA with a specific 5’ end modification, we prepared a pair of strands: sense and antisense, both with the same 5’ end modification, and annealed it (Figure 1A). By using different cap analogs for the *in vitro* transcription, we obtained dsRNAs with cap0, cap1, or cap2 structure at the 5’ end. Furthermore, our goal was to generate dsRNA of sufficient length to be recognized not only by RIG-I, but also by host receptors whose mode of action depends primarily on the recognition of RNA duplex. The optimal length to activate OAS proteins is more than 50 bp;^19,26,39^ for the PKR substrate, it should be at least 30 bp long.^15^ For MDA5, the dsRNA should be as long as 1 kbp,^23^ although other reports have shown that this host sensor is able to recognize much shorter transcripts than 1 kbp.^24,40^ However, because the exact mechanism of dsRNA selection by MDA5, as well as its activation, has not been fully characterized, we decided to prepare a blunt-ended dsRNA 552 bp in length (Figure 1B and S1A). To ensure that the used transcripts were of high purity and free from T7 RNA polymerase by-products, such as dsRNA impurities and ssRNA molecules of unintended size, we performed high performance liquid chromatography (HPLC) purification of the *in vitro* transcribed RNA. Finally, to obtain homogeneous capped transcripts, HPLC-purified RNAs were enzymatically treated to remove the uncapped versions. Once the sense and antisense strands used for dsRNA formation were purified, they were heated and cooled slowly to form duplexes. The efficiency of duplex formation was verified by agarose gel electrophoresis (Figure 1C and S1B).

**Figure 1.**
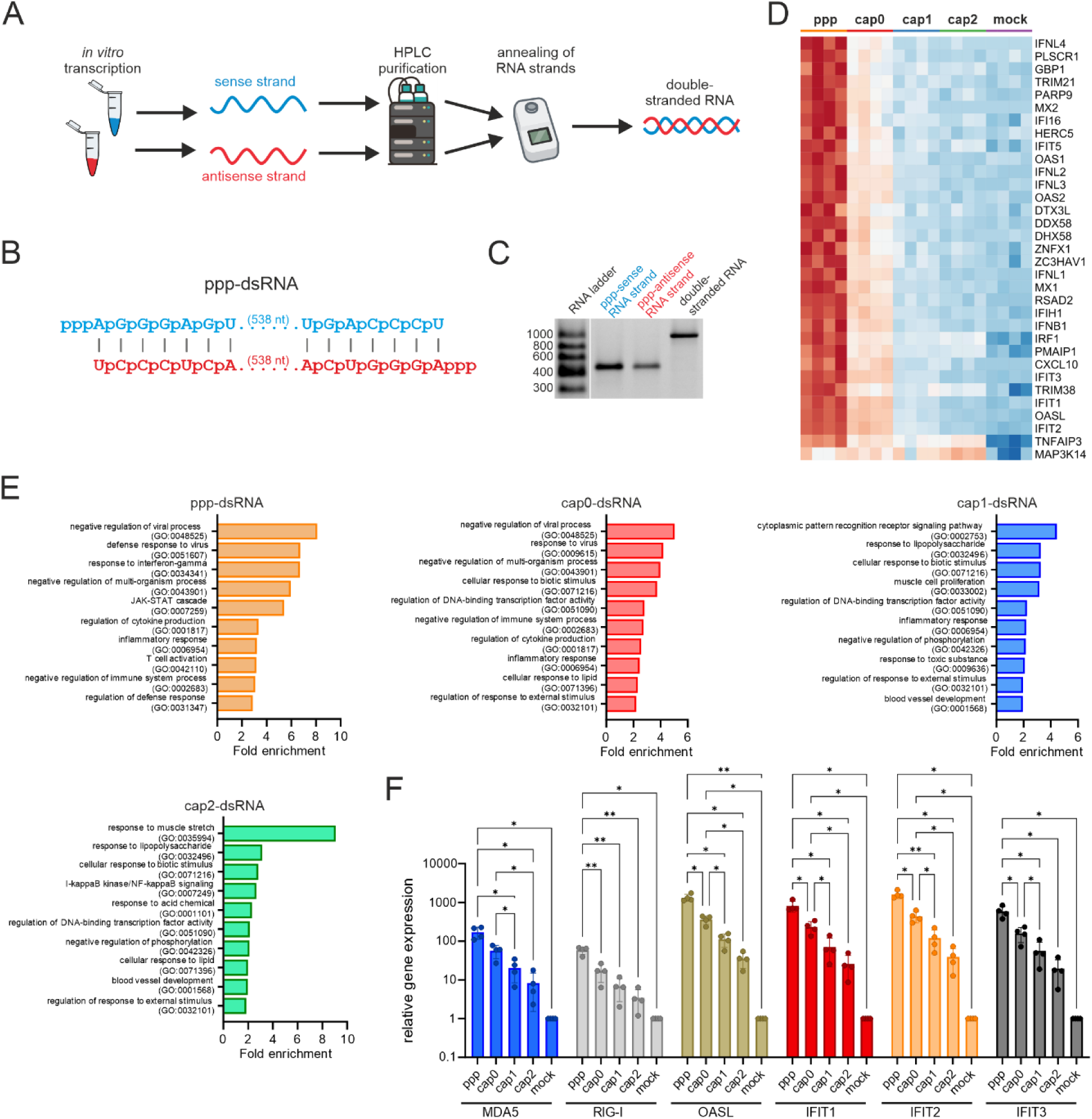
2’-*O*-methylation of cap structure severely compromises double-stranded RNA (dsRNA) immunogenicity. (A) Scheme of generation of dsRNA. First, two strands used for dsRNA formation are obtained by separate *in vitro* transcription reactions with T7 RNA polymerase. Both resulting transcripts are then purified by high performance liquid chromatography (HPLC). Capped versions of the HPLC-purified single stranded RNAs (ssRNAs) are enzymatically treated with 5’ polyphosphatase and XRN1 to eliminate the uncapped fraction of transcripts (this step is not shown in the scheme). Finally, a pair of complementary strands is heated and slowly cooled down to force duplex formation, *i.e.* dsRNA. (B) Schematic representation of *in vitro* transcribed dsRNA with triphosphate group. Other dsRNAs with differently modified 5’ ends generated for this study are shown in Figure S1A. (C) Analysis in agarose gel of dsRNA forming strands: the sense and antisense one with a triphosphate group at the 5’ end. On the far right, complementary strands after annealing. Agarose gel analysis of other dsRNA with differently modified 5’ ends generated for this study are shown in Figure S1B (D) Heatmap displaying differentially expressed genes in A549 cells transfected with variously 5’ end modified dsRNA (ppp-, cap0-, cap1-, and cap2-dsRNA). This representation is limited only to genes assigned to “defense response to virus” gene ontology (GO) term (GO:0051607). Full heatmap of 0.4% upregulated genes for all analyzed samples is presented in Figure S3 (E) Top ten overrepresented GO terms for A549 cells transfected with each of tested dsRNA. (F) Validation of RNA-Seq data by RT-qPCR – changes in gene expression upon cell transfection with differently capped dsRNA. Bars represent the mean value of mRNA level change (relative gene expression) ± SD from four independent biological replicates, each independent biological replicate consists of a single transfection reaction. Each point represents data from one independent biological replicate. Data were normalized to mock treated cells. Statistical significance: * P < 0.05, ** P < 0.01 (one-way ANOVA with Turkey’s multiple comparisons test). Only statistically significant differences were marked on the graph.

With the set of 5’ end modified dsRNA in our hands we tested whether the presence of cap structure indeed affects dsRNA immunogenicity. To this end, we transfected human A549 lung epithelial cells with dsRNA carrying cap1 or triphosphate group at its 5’ end. After 5 h, as we expected, the level of interferon β (IFNβ) transcript was much higher in cells transfected with ppp-dsRNA than those in cells transfected with its counterpart bearing cap1 (Figure S2). This result encouraged us to perform genome-wide transcriptomic analysis to precisely define the contribution of each of 5’ end modifications to immunogenicity of dsRNA. Analogically, RNA sequencing (RNA-Seq) revealed that the most immunogenic version of the dsRNA was the one with a triphosphate group at the 5’ end, which was reflected in the highest number of upregulated genes in all the samples analyzed (Figure 1D and S3). Gene Ontology (GO) analysis showed that the group of genes associated with the immune and antiviral pathways was the most upregulated (Figure 1E). The presence of the cap structure reduces the immunogenicity of dsRNAs. However, unexpectedly, the incorporation of 2’-*O*-methylation within the cap structure (cap1) almost completely abolished dsRNA immunogenicity (Figure 1D and S3). The presence of cap0, and therefore, the absence of 2’-*O*-methylation, results in a moderate immunostimulatory effect of dsRNA on A549 cells. In contrast, the addition of a second 2’-*O*-methylation at the 5’ end of the dsRNA (cap2) only slightly altered the gene expression profile observed for cap1-dsRNA-transfected cells, indicating that the presence of only one 2’-*O*-methylation is fully sufficient to severely reduce the immunogenic potential of the dsRNA (Figure 1D, S3, and S4). In addition, GO terms associated with innate immune or antiviral responses were no longer the most enriched in cells transfected with cap1- or cap2-dsRNA (Figure 1E). The RNA-Seq results were confirmed by independent analysis of the expression levels of selected ISGs (IFIT1, IFIT2, IFIT3, MDA5, OASL, and RIG-I) by RT-qPCR (Figure 1F). These results strongly suggest that the 5’ end of dsRNA is the primary trigger for cellular inflammation. The effect of the cap presence at the 5’ end of dsRNA on its immunogenicity is consistent with our knowledge about RIG-I activation: 2’-*O*-methylation of the cap structure significantly reduces RIG-I affinity for dsRNA.^20^ Similarly, it is thought that 2’-*O*-methylation in the cap structure may also contribute to the evasion of dsRNA recognition by MDA5,^12,25^ but other reports indicate that the length of the duplex in the transcript is the main activator of this receptor.^23,41^ Nevertheless, the dsRNA we prepared was able to evade recognition by the RLR pathways due to the 2’-*O*-methylation incorporated into its cap structure. Therefore, this tool allowed us to study the activation of cell growth inhibitory pathways independently of the cell’s inflammatory response.

### Despite reduced immunogenicity, capped dsRNAs are still detected by PKR

We hypothesized this indicates that dsRNAs with significantly reduced immunogenicity are no longer recognized by host receptors. Alternatively, dsRNAs are recognized by host receptors despite their lack of immunogenicity, and host cells can take countermeasures to eliminate the potential threat of dsRNAs. To answer these questions, we performed a dsRNA interactome capture experiment to determine whether modifications at the 5’ end of dsRNA affect their binding partners in living cells. Importantly, our goal was to identify proteins that are involved in dsRNA recognition as soon as they appear in the cytoplasm of the host cell and not proteins that are ISG products that play a role in cellular defense only after dsRNA has been successfully recognized by the cell’s innate immunity. Therefore, we first tested the duration of transfection to select appropriate conditions for studying the primary cellular response at the protein level. We verified that 5 h after transfection, there was no increase in the level of ISGs products with highly immunogenic ppp-dsRNA, in contrast to 24 h after transfection (Figure S5). Therefore, A549 cells were transfected with a set of dsRNAs with different 5’ end modifications for 5 h. Moreover, to identify host proteins that bind to dsRNA as accurately as possible, we used crosslinked proteins and their associated nucleic acids, by exposing plated cells to ultraviolet (UV) light. After cell lysis, only ribonucleoprotein (RNP)complexes formed on the dsRNA were fished out using dsRNA-specific antibodies. The captured RNP complexes were subjected to semi-quantitative tandem mass spectrometry (Figure 2A). To ensure that the analyzed cells responded in the expected manner to differently capped dsRNAs, we tested the expression levels of three ISGs (IFIT1, MDA5, and RIG-I) in cell lysates (Figure 2C). Proteomic analysis revealed that the more immunogenic the dsRNAs, the more proteins were bound to them (Figure 2B, S6, and S7). In addition, GO analysis showed that among the proteins enriched in immunoprecipitates (IPs) are factors involved in the innate immune response, but their number decreased when 5’ capped dsRNA was delivered to the cells (Figure S7). However, regardless of the modification of the 5’ end of dsRNA, several proteins were always present in IPs, including factors involved in the innate immune response, namely adenosine deaminases acting on RNA (ADAR) and PKR (EIF2AK2) (Figure 2B, S6, and S7). ADAR is responsible for adenosine-to-inosine editing within endogenous transcripts in mammalian cells and plays a critical role in the regulation of innate immune activation by suppressing endogenous RNA sensing.^42^ However, ADAR activity on foreign dsRNA can have pro-viral effects.^43,44^ In addition, ADAR activity is unlikely to lead to global changes in host cell metabolism, which may help host cells rearrange their metabolism in response to viral threats. PKR, in contrast, is a dsRNA-binding kinase that phosphorylates the translation initiation factor eIF2α in response to various stress stimuli.^16^ Importantly, PKR is one of the key sensors responsible for detecting foreign RNAs during viral infection.^2,9^ PKR activation leads to the shutdown of global host cell translation to minimize viral spread. Thus, PKR appears to be a protein that can recognize dsRNA regardless of its immunogenicity and adjust the cell’s metabolism to the emerging threat.

**Figure 2.**
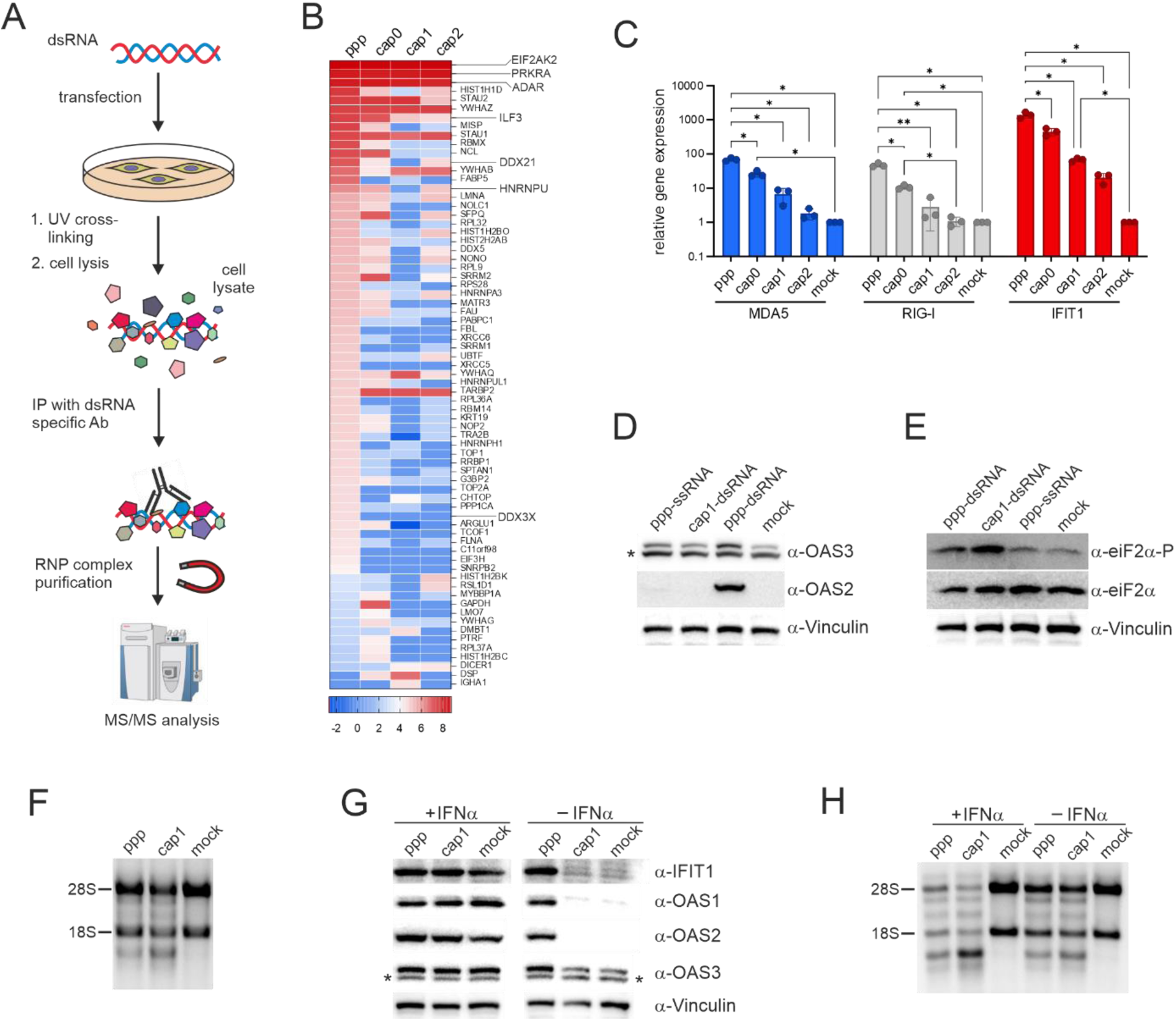
Less immunogenic double-stranded RNA (dsRNA) is still recognized by the PKR and OAS/RNase L pathways. (A) Scheme of proteomics approach used in this study. After transfection with differentially capped dsRNA for 5 h, A549 cells are cross-linked with ultraviolet (UV) light and lysed. Ribonucleoprotein (RNP) complexes containing dsRNA are fished out from the cell lysates using dsRNA-specific antibodies. The composition of the captured RNP complexes is analyzed by tandem mass spectrometry (MS/MS). (B) Protein specificity heatmap. This heatmap visualizes the specificity of identified proteins across different experimental conditions. The Specificity value is calculated as the log2 intensity ratio compared to the control. Proteins with a Specificity value greater than 4.0 in at least one of the conditions are shown. Note that this heatmap is an excerpt from the full heatmap shown in Figure S6. (C) Verification of the activation of the RLR pathways by RT-qPCR in A549 cell lysates after transfection with dsRNAs bearing different modifications at the 5’ end for 5 h. Bars represent the mean value of mRNA level change (relative gene expression) ± SD from three independent biological replicates. Each point represents data from one independent biological replicate. Data were normalized to mock treated cells. Statistical significance: * P < 0.05, ** P < 0.01 (one-way ANOVA with Turkey’s multiple comparisons test). Only statistically significant differences were marked on the graph. (D, E) Triphosphorylated ssRNA (ppp-ssRNA) does not activate RLR and PKR pathways. A549 cells are transfected with ppp-dsRNA, cap1-dsRNA, or ppp-ssRNA for 24 h and (D) ISG products expression levels as well as (E) phosphorylation status of eIF2α are assessed by western blotting. * indicates unspecific band. (F) RNase L activity in A549 cells assessed by ribosomal RNA (rRNA) integrity. Total RNA was isolated after 24 h transfection with ppp-and cap1-dsRNA and analyzed in 1x TBE agarose gel. (G) Activation of RLR pathways in interferon-treated (+IFNα) (200 U/ml) and untreated (-IFNα) cells assessed by expression of ISGs products. A549 cells were transfected with ppp- and cap1-dsRNA for 24 h. * indicates unspecific band. (H) RNase L activity is dependent on the presence of dsRNA and independent of the immune state of the cell. RNase L activity in A549 cells assessed by rRNA integrity. Total RNA was isolated after 24 h pretreatment with INF α (200 U/ml), followed by 24 h transfection with ppp- or cap1- dsRNA, and analyzed in 1 x TBE agarose gel.

We then tested whether dsRNA with reduced immunogenicity is recognized by PKR and whether this cell receptor, upon detection of cap1-dsRNA, can trigger a signaling cascade leading to global translation repression. We also investigated whether the lack of cellular inflammatory activation affects the global shutdown of host protein biosynthesis by PKR. To this end, we examined the phosphorylation status of eIF2α in cells transfected with the highly immunogenic ppp-dsRNA and a dsRNA with significantly reduced immunogenicity – cap1-dsRNA. First, we tested whether we could observe the activation of innate immunity due to the presence of immunogenic dsRNA. As expected, we observed increased levels of ISG products (IFIT1, OAS1, OAS2, and OAS3) in cells transfected with ppp-dsRNA, but not with cap1-dsRNA for 24 h (Figure 2D). However, elevated levels of phosphorylated eIF2α (p-eIF2α) were observed regardless of the dsRNA used for cell transfection (Figure 2E). Furthermore, we were wondering whether PKR activation and subsequent phosphorylation of eIF2α occurs whenever foreign RNA is present in the cell cytoplasm. Mayo and Cole suggested that PKR can bind not only to dsRNA, but also to ssRNA, albeit with lower affinity.^45^ Nevertheless, recognition of ssRNA by PKR also leads to its activation *in vitro*. We therefore tested whether PKR activation is based solely on recognition of the structure of RNA duplex, or whether the presence of exogenous ssRNA is sufficient to lead to eIF2α phosphorylation. We have previously shown that transfection of mammalian cells with exogenous single stranded transcripts, even those with triphosphate groups at their 5’ ends, did not lead to stimulation of the antiviral and interferon responses.^36^ Our western blotting analysis showed exactly the same phenotype, we observed no increase in expression of ISG products upon transfection of A549 cells with triphosphorylated-ssRNA (ppp-ssRNA) (Figure 2D). Finally, we showed that when cells were transfected with ssRNA bearing triphosphate group at its 5’ end, the level of p-eIF2α remained unchanged compared to the mock control (Figure 2E).

Taken together, we showed that the double-strandedness is an essential feature of exogenous transcripts for the activation of the PKR-dependent innate immune response pathway. Furthermore, this activation does not depend on either the immunogenic potential of the dsRNA or the activation of cellular inflammation.

### OAS/RNase L pathway does not depend on activation of cellular inflammation

The host cell adapts to potential dsRNA threats not only by slowing down global translation, but also by eliminating ssRNAs which can be a source of dangerous dsRNAs, for example during viral replication.^2,9^ The aforementioned ssRNA degradation is achieved primarily through activation of OAS/RNase L pathway, in which OAS proteins sense dsRNAs and then produce 2′-5′-linked oligoadenylates (2-5As), that ultimately serve as activators of RNase L.^19^ Induction of this mechanism leads to degradation not only of the virus ssRNAs but also of the host molecules. Although OASs belong to the ISGs, *i.e.* their levels are greatly increased during viral infection, it has been suggested that the basal level of OAS proteins may be sufficient to activate RNase L even before the onset of inflammation.^32^ Thus, it appears that RNase L can be activated regardless of the immune state of the host cell, but only if OASs produce enough 2-5As as activators of this endonuclease. Intriguingly, other reports have shown that, among the three OAS proteins, OAS3, which has the highest affinity for dsRNA^26,27^ and RNase L activation, depends primarily on its expression during viral infection.^27^

Despite the fact that none of the oligoadenylate synthetases were identified in our proteomic analysis, the OAS3 protein seems to be a suitable candidate for a factor capable of responding effectively to dsRNA regardless of the immune state of the host cell. Interestingly, our analysis of RLR pathway activation showed that the basal level of OAS3 was relatively high and moderately increased in A549 cells after interferon treatment (Figure 2G). In contrast to OAS3, basal expression levels of OAS1 and OAS2 were limited. Considering this, we examined RNase L activity in A549 cells transfected with dsRNAs differing in immunogenic potential by checking the integrity of the 18S/28S rRNA isolated from these cells. We observed that 18S and 28S rRNA were degraded regardless of the 5’ end modification (Figure 2F), suggesting that activation of cellular inflammation is not necessary for OAS proteins to recognize dsRNAs and activate RNase L. Presumably, the observed phenomenon is caused by OAS3, since OAS1 and OAS2 levels are very low and unchanged in cells transfected with cap1-dsRNA (Figure 2D,E). Finally, as expected, treatment of A549 cells with IFNα, although leading to a high upregulation of all three OAS proteins (Figure 2G), did not result in the appearance of rRNA fragmentation in mock-treated cells (Figure 2H), meaning that only the presence of dsRNA in the cytoplasm of the cell activates the OAS/RNase L pathway. Importantly, this activation is independent of the immune state of host cells.

### Epitranscriptomic marks do not shield dsRNA from recognition by RLR and PKR pathways

In addition to adding cap structures to their transcripts,^28^ viruses install epitranscriptomic marks within RNA to generate RNAs that resemble human transcripts as closely as possible.^29^ Importantly, internal modifications found in viral RNA are the same as those present in the human transcriptome. Therefore, we decided to test whether the presence of any of the three most prevalent post-transcriptional modifications in the human RNA, *i.e.* m^6^A, Ψ, or m^5^C, could affect dsRNA immunogenicity.^29^ To study this, we prepared a set of dsRNAs by *in vitro* transcription in which each particular nucleotide was replaced with its modified version (Figure S8A). In other words, transcripts with m^6^A had all the adenosines in both dsRNA strands changed to their *N*6-methylated analogs. RNAs carrying Ψ and m^5^C were obtained in an analogous manner, *i.e.* in the sequence of the former transcripts all uridines were replaced by pseudouridines, and for the latter all cytidines were changed to their 5-methyl counterparts. In addition, dsRNAs with modifications were obtained in two versions, with triphosphate or cap1 at the 5’ ends of each dsRNA strand. Regardless of the chemical modifications introduced within the RNA sequence, in all cases each capped transcript has an adenosine with a 2’-*O*-methyl group at the position of the first transcribed nucleotide. Therefore, we used pppApG dinucleotide analog as the initiator of the *in vitro* transcription reaction to prepare triphosphorylated RNA containing internal m^6^A, which allowed us to obtain RNA starting with adenosine instead of *N*6-methyladenosine (Figure S8A). We verified the efficiency of modified dsRNA formation using an agarose gel (Figure S8B). In addition, the stability of RNA duplexes was tested using RNase I (Figure S8C), which specifically degrades single-stranded transcripts and preserves double-stranded molecules.^46^ The prepared dsRNAs were used to study the activation of the interferon pathway by introducing differently modified dsRNA into A549 cells which transiently expressed the firefly luciferase gene under the control of the INFβ promoter.

We found that regardless of the chemical modification introduced within ppp-dsRNA, it was immunogenic to the same extent as its unmodified counterpart (Figure 3A). Interestingly, the modified cap1-dsRNAs were slightly less immunogenic than the unmodified capped variants, but these differences were not statistically significant. Furthermore, at the protein level, we observed no differences in the expression levels of selected ISGs (IFIT1, OAS1, OAS2, and OAS3) between the unmodified dsRNAs and their modified counterparts (Figure 3B and S9A). Taken together, it seems that the only feature of dsRNA that affects its immunogenic potential and leads to the activation of RLR pathways is the presence of cap structure at transcript 5’ ends.

**Figure 3.**
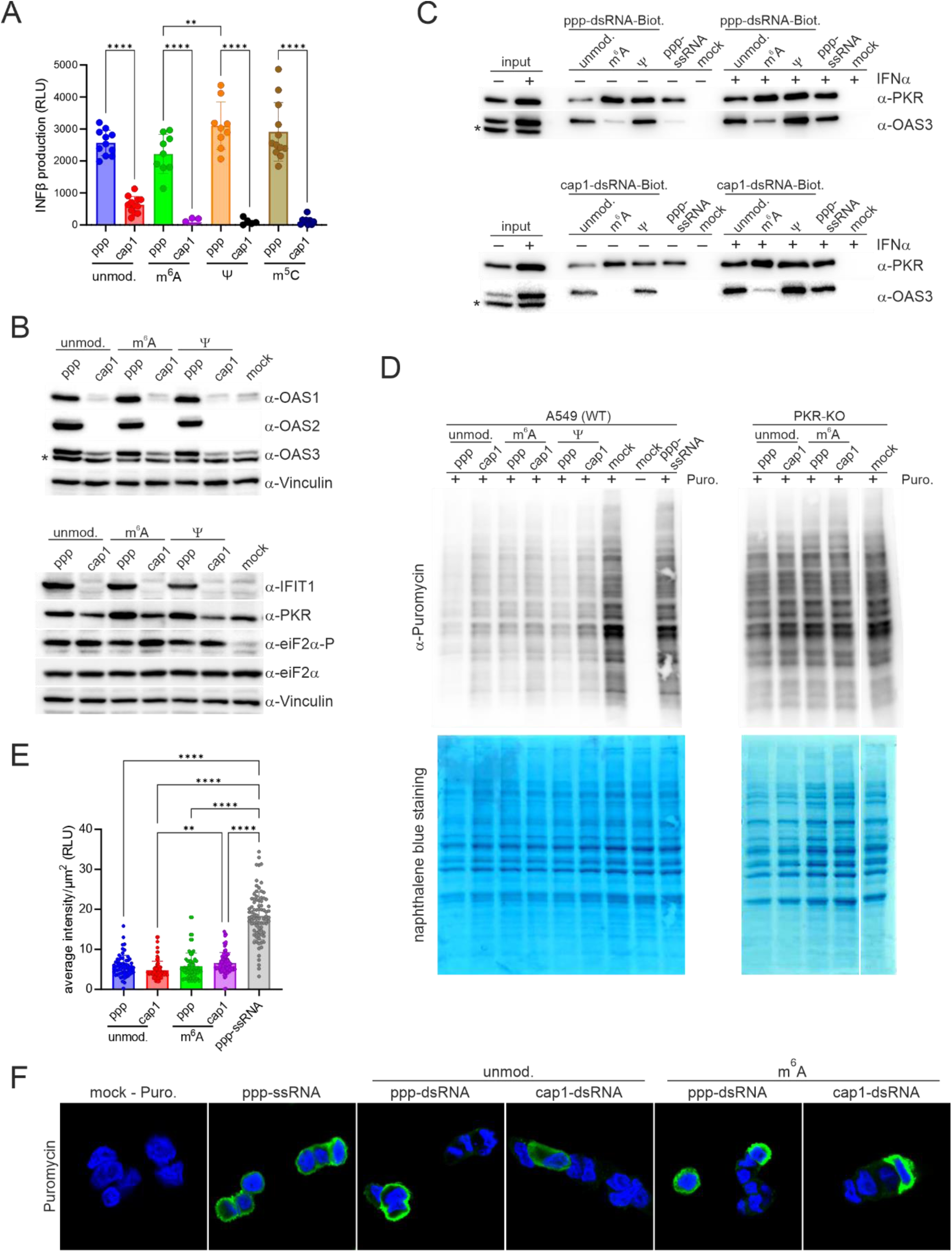
Presence of *N*6-methyladenosine (m^6^A), pseudouridine (Ψ), and 5-methylcytosine (m^5^C) within double-stranded RNA (dsRNA) does not contribute to the reduction of dsRNA immunogenicity, nor affects dsRNA recognition by PKR. (A,B) Activation of RLR and PKR pathways is not affected by the presence of post-transcriptional modifications within dsRNA. (A) Interferon β (IFNβ) production is assessed by measuring the luminescence of A549 cells transiently transfected with a plasmid encoding firefly luciferase under the control of the IFNβ promoter. These cells are transfected with dsRNA for 24 h and then firefly luciferase activity is measured. Bars represent the mean value ± SD from at least three independent biological replicates (each biological replicate consist of three technical replicate). Each point represents data from one technical replicate. Statistical significance: ** P < 0.01, **** P < 0.0001 (one-way ANOVA with Turkey’s multiple comparisons test). Only statistically significant differences were marked on the graph. (B) A549 cells are transfected with triphosphorylated- (ppp-) and cap1- dsRNA carrying different epitranscriptomic marks for 24 h and ISG products expression levels as well as phosphorylation status of eIF2α are assessed by western blotting. * indicates unspecific band. (C) Co-purification of endogenous proteins from lysates of interferon α (IFNα)-treated (200 U/mL) and untreated A549 cells with biotinylated ppp- or cap1-dsRNA. PKR and OAS3 are detected in precipitates by western blotting. This is a cropped version of Figure S10. * indicates unspecific band. (D-F) The levels of newly synthetized proteins are assessed by puromycin incorporation assay. A549 and PKR-KO cells are transfected with ppp- and cap1- dsRNA carrying different post-transcriptional modifications for 24 h and then puromycin (1 µg/ml) is added for 30 min. (D) The levels of newly synthetized proteins are assessed by western blotting with anti-puromycin antibodies. Naphthalene blue staining is performed to control the loading. Verification of PKR knocked out (PKR-KO) cells is shown in Figure S11. (E-F) The levels of newly synthetized proteins are assessed by immunofluorescence with anti-puromycin antibodies. (E) Average intensity of puromycin staining in cells is presented in (F) (between 81 and 173 cells are analyzed for each transfection). Bars represent the mean value ± SD from all analyzed cells. Each point represents data from one cell. Statistical significance: ** P < 0.01, **** P < 0.0001 (one-way ANOVA with Turkey’s multiple comparisons test). Only statistically significant differences were marked on the graph.

It is known that post-transcriptional modifications can affect the recognition of dsRNA by host factors that sense dsRNA based on its double-strandedness and not on the modifications present at its 5’ ends.^2,9,29^ PKR is thought to be one of such sensors.^47^ It has been previously reported that transfer RNAs (tRNAs) lacking post-transcriptional modifications can activate PKR *in vitro* and *in cellulo*.^48,49^ To investigate if and how the three most abundant post-transcriptional modifications in the human transcriptome affect PKR binding to dsRNA, we performed pull down experiments. We fished out proteins from the lysates of interferon-treated and untreated A549 cells using biotinylated dsRNA containing selected chemical modifications (Figure 3C). Since none of the tested post-transcriptional modifications affects the immunogenicity of dsRNA (Figure 3A,B), we used only two of them, m^6^A and Ψ, to prepare biotinylated dsRNA for the pull-down experiment. PKR binds to each dsRNA regardless of its modification. Similarly, 5’ end cap did not affect dsRNA recognition by PKR. PKR is known to bind both dsRNA and ssRNA,^45^ and as expected, we found that it also bound to ssRNA in our analysis. These observations are consistent with the eIF2α phosphorylation status in A549 cells studied after transfection with dsRNA carrying m^6^A or Ψ (Figure 3B and S9A). None of the tested modifications affected phosphorylation of the inspected protein.

We further investigated the hypothesis that post-transcriptional modifications do not prevent inhibition of global protein biosynthesis *via* a PKR-dependent pathway. We studied the levels of the newly synthesized puromycin-labeled proteins in A549 cells transfected with modified or unmodified dsRNAs (Figure 3D). In western blotting analysis protein levels with incorporated puromycin was found to decrease in the presence of epitranscriptomic marks and 5’ end cap in dsRNAs. An analogous conclusion was drawn from the immunofluorescence studies of puromycin-labeled proteins in A549 cells (Figure 3E,F). Consistent with the assumption that PKR is the host factor primarily responsible for slowing the rate of protein biosynthesis in virus-infected cells, we observed that regardless of the dsRNA used for transfection, no changes in translation were detected in PKR-KO cells (Figure 3D).

Altogether, obtained results lead to the conclusion that the presence of the three most widespread post-transcriptional modifications in the human transcriptome, *i.e.* m^6^A, Ψ, and m^5^C, within dsRNA does not contribute to the reduction of dsRNA immunogenicity, nor affects dsRNA recognition by PKR and subsequent translational repression.

### *N*6-methylation of adenosine shields dsRNA from being recognized by the OAS/RNase L pathway

A pull-down experiment with biotinylated dsRNA showed that one of the tested epitranscriptomic marks abolished OAS3 binding to dsRNA (Figure 3C), in contrast to the unaffected interactions between RNA duplexes and PKR. As expected, this was observed independently of the 5’ end modification of the dsRNA. Whether the presence of m^6^A affects dsRNA recognition by other OAS proteins could not be confirmed in our experimental setup (Figure S10). However, the absence of OAS1 and OAS2 in IPs can be explained by the fact that these two proteins have a much lower affinity for dsRNA than OAS3.^26,27^ To verify whether the *in vitro* observation that the presence of m^6^A affects OAS3 binding to dsRNA translates into a phenotype in living cells, we examined the integrity of rRNAs after transfection of A549 cells with various modified dsRNAs. We found that the presence of m^6^A protected dsRNA from recognition by OAS proteins *in cellulo* (Figure 4 and Figure S9B). Importantly, the delivery of ppp-dsRNA containing m^6^A did not lead to the activation of RNase L in A549 cells. This observation supports the assumption that activation of cellular inflammation is dispensable for effective dsRNA recognition by OAS proteins, at least for OAS3 and activation of RNase L. Furthermore, in MAVS-KO cells, *i.e.* cells with disabled RLR pathways, we observed degradation of rRNAs unless these KO cells were transfected with dsRNA carrying the m^6^A modification (Figure 4). Only the elimination of RNase L completely inhibited rRNA degradation, regardless of the dsRNA used. Similar to that in RNase L-KO cells, a significant decrease in rRNA degradation was observed in OAS3-KO cells. However, some rRNA degradation can still be observed in OAS3-KO cells upon delivery of immunogenic ppp-dsRNA, but this is probably due to the increased expression of OAS1 and OAS2 and activation of RLR pathways. However, in the case of ppp-dsRNA with m^6^A, no rRNA degradation was observed, strongly suggesting that this chemical modification protected dsRNA not only from recognition by OAS3, but also from detection by OAS1 and OAS2.

**Figure 4.**
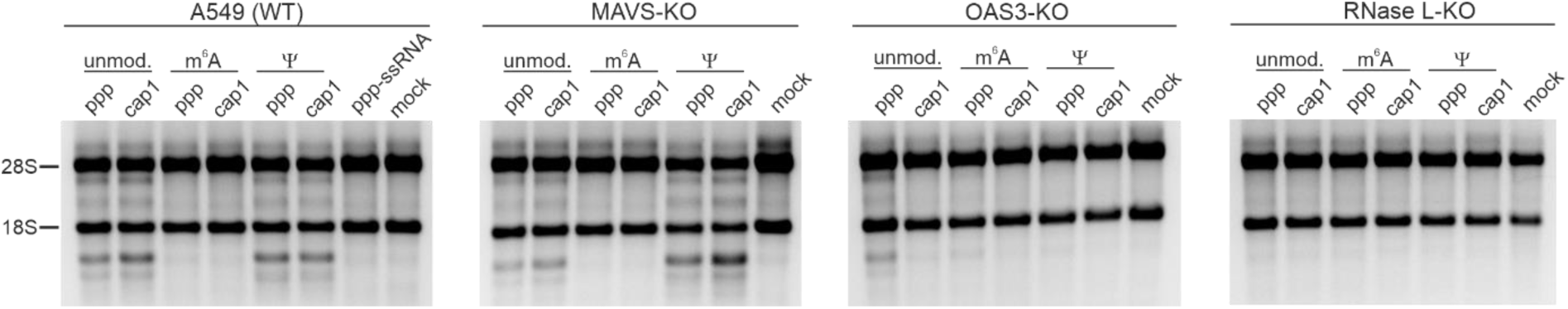
The presence of m^6^A within double-stranded RNA (dsRNA) affects RNase L activity. RNase L activity in A549, MAVS-KO, OAS3-KO, and RNase L-KO cells assessed by ribosomal RNA (rRNA) integrity. Total RNA is isolated after 24 h transfection with post-transcriptionally modified (m^6^A or Ψ) ppp- or cap1- dsRNA and analyzed in 1x TBE agarose gel. Verification of MAVS, OAS3, and RNase L knocked out in MAVS-KO, OAS3-KO, and RNase L-KO cells, respectively, was shown in Figure S11.

### PKR clustering on dsRNAs occurs regardless of the immune state of the cell

First, to repress translation, PKR is activated by autophosphorylation upon binding to dsRNAs and becomes dimerized.^50–53^ Only as a dimer PKR is capable of phosphorylating eIF2α, which subsequently leads to translational shutdown.^54,55^ In addition, Zappa et al. recently showed that PKR in human cells transfected with poly(I:C) forms not only dimers, but also higher order assemblies.^56^ The exact reason PKR forms clusters is not yet clear, but it has been proposed that PKR clustering may fine-tune its activity; translational inhibition is moderately reduced by PKR clustering.^56^ Disruption of PKR clusters *in cellulo* leads to a moderate increase in eIF2α phosphorylation. Because we already knew that post-transcriptionally modified dsRNA could be effectively recognized by PKR (Figure 3C-F), we wondered whether the presence of epitranscriptomic marks could fine-tune PKR activity through clustering. Furthermore, we were interested in whether the immunogenic potential of dsRNA, and thus the immune state of the host cells, could influence this phenomenon. We prepared a set of Cy5-labeled triphosphorylated and capped versions of dsRNAs. Because none of the chemical nucleotide modifications tested affected PKR binding or translation repression, we examined the clustering phenomenon only for dsRNAs with m^6^A and their unmodified counterparts. Upon transfection of fluorescent dsRNAs into A549 cells, we observed spots in the cell cytoplasm containing introduced transcripts (Figure 5A), as previously reported.^30,35,56^ In addition, we found that the majority of foci with Cy5-labeled dsRNA (from 60% to 81%) colocalized with PKR, regardless of the modification at the 5’ end of the transcript (Figure 5B). The colocalization of PKR with dsRNA was also not affected by the presence of m^6^A in dsRNA. Importantly, almost all signals recorded for PKR (from 84% to 97%) overlapped with those observed for dsRNA. The higher percentage of PKR colocalized with dsRNA than dsRNA colocalized with PKR suggests that dsRNA first forms cytoplasmic foci, and then PKR is attracted to these foci and begins to form clusters. This hypothesis is supported by the fact that PKR is not required for the formation of dsRNA condensates. The dsRNA foci were still visible in PKR-KO cells transfected with Cy5-labeled dsRNA (Figure S12A,B). Our observation is consistent with that of a recent study by Corbet et al. showing that the presence of PKR is dispensable for the formation of poly(I:C) foci.^30^

**Figure 5.**
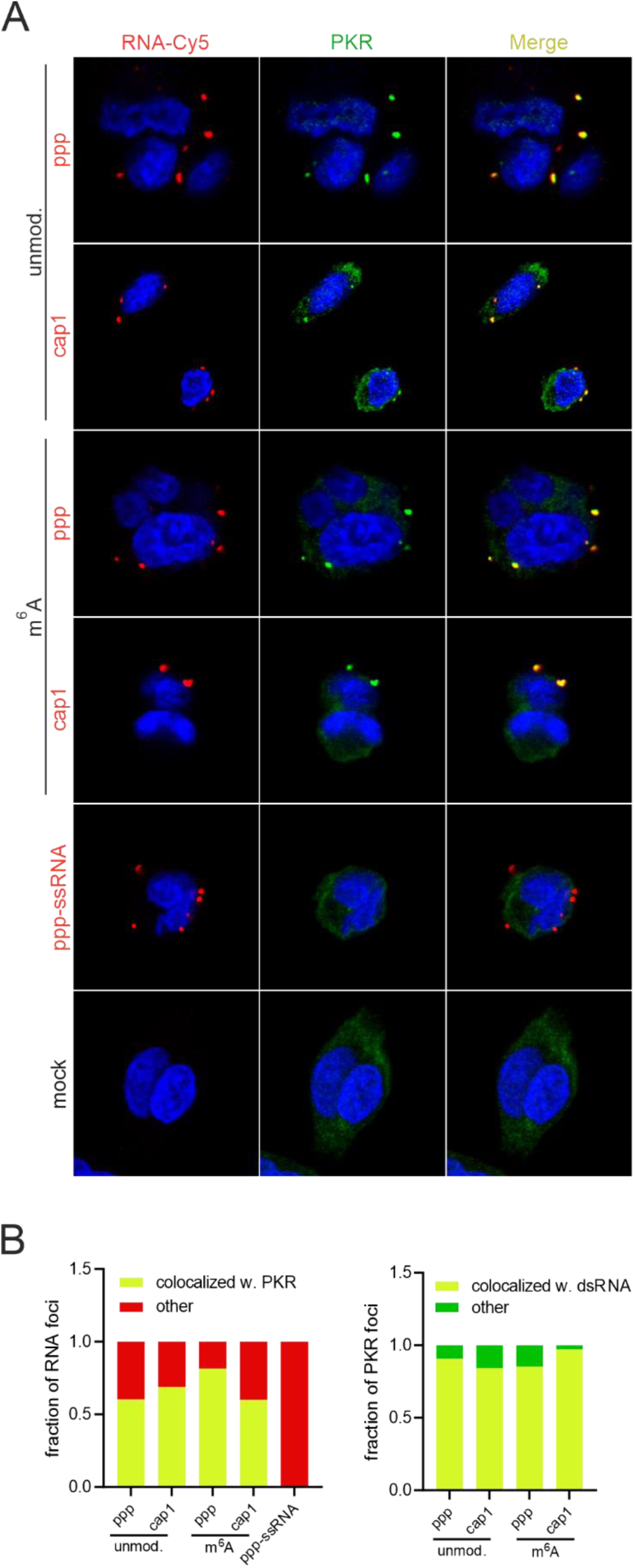
PKR clusters on differently modified double-stranded RNA (dsRNA). (A) Immunofluorescence analysis of PKR and Cy5-labeled RNA in A549 cells. Cells are transfected either with Cy5-labeled ppp- or cap1-dsRNA bearing m^6^A or with unmodified counterparts or with triphosphorylated ssRNA (ppp-ssRNA) also labeled with Cy5. (B) Quantification of (left) Cy5-labeled RNA colocalization with PKR or (right) PKR colocalization with Cy5- labeled transcripts (between 127 and 874 cells are analyzed for each transfection). Each bar represents all foci counted, yellow color represents the fraction of foci in which RNA colocalized with PKR, red and green colors represent the fraction of foci in which only signal from PKR and RNA was observed, respectively.

Interestingly, RNA condensates were also present in cells transfected with ssRNA (Figure 5A). However, this observation is not unexpected, as the occurrence of punctate signal containing mRNAs in mammalian cells transfected by lipofection is already well documented.^57–62^ The aim of PKR clustering is to limit its activity,^56^ namely to reduce eIF2α phosphorylation, and it is possible that also foci containing ssRNA may also form clusters with PKR. However, in this case, PKR did not co-localize with the RNA condensates (Figure 5A,B). Thus, the lack of colocalization of PKR with ssRNA provides further evidence that despite its affinity for ssRNA *in vitro*,^45^ this kinase specifically recognizes only dsRNA in living cells. Furthermore, the observation that both ssRNA and dsRNA form RNA condensates indicates that the presence of these foci is not due to the presence of specific nucleic acids in the cell cytoplasm. In all these cases, the transcripts were introduced into cells by means of lipofection, which is based on the formation of complexes, *i.e.* lipoplexes, between nucleic acids and positively charged lipids present in the transfection reagent.^57,63^ Thus, the spots observed in the cells are only lipoplexes from which the cargo (nucleic acids) has not yet been released.^63–65^ This clearly demonstrates that the formation of RNA condensate is independent of PKR action and that this phenomenon probably occurs independently of the action of any other host cell protein. Moreover, the presence of RNA condensates formed from foreign transcripts in the cell is simply due to the way in which nucleic acids are introduced into the cell.

## Discussion

Our studies have shown that effective dsRNA recognition in mammalian cells is achieved independent of RLR pathways activation. This process is mediated by the action of host cell sensors involved in cell growth inhibition such as OASs and PKR. Although they are ISGs, their basal levels are sufficiently high to mount action against dsRNA without prior induction of inflammation. As we did not observe an increase in the expression of ISG products in cells with activated OAS/RNase L and PKR pathways, we postulate that the activation of these defense mechanisms does not contribute to cellular inflammatory processes, or that this contribution is low. Activation of cellular inflammation was observed for dsRNA with a triphosphate group at the 5’ ends and to some extent in cells treated with dsRNA carrying a cap0 structure. In contrast, the presence of a 2’-*O*-methylated version of the cap structure almost completely abolished dsRNA immunogenicity. We observed only a slight upregulation of some gene expression in cells treated with cap1- or cap2-dsRNA, but at the protein level the expression of ISGs products for cells transfected with dsRNA with 2’-*O*-methylated cap structure was at the mock control level.

Among the RLRs, two of them, namely RIG-I and MDA5, are directly involved in dsRNA recognition and activation of the signaling cascade leading to the expression of ISGs, cytokines, and interferons.^11–13,21^ It is well documented that RIG-I recognizes dsRNA based on its 5’ end structure and has the highest affinity for triphosphorylated dsRNA, but only the presence of a 2’-*O*-methyl group within the cap structure eliminates RIG-I’s interaction with dsRNA.^2,11,20–22^ This is consistent with our observations that only the presence of a 2’-*O*-methylated cap structure reduces the immunogenic potential of dsRNA. However, our data showed that transfection of cells with dsRNA containing only cap0 resulted in a significant decrease in ISG expression. Further modification of the 5’ end by introducing 2’-*O*-methyl groups into the cap structure led to a significant decrease in immunogenicity, which was particularly striking at the protein level, and there was no difference in the level of ISGs products in cells transfected with cap1-dsRNA compared to mock controls. The lack of immunogenicity of cap1- and cap2-dsRNAs raises questions about the role of MDA5 in dsRNA detection. MDA5, unlike RIG-I, is not likely to recognize dsRNA based on modifications of its 5’ ends, but is rather activated by the long duplex structure.^23,24^ It is thought that the optimal MDA5 substrate should be longer than 1 kbp,^23^ but there are also reports that MDA5 is able to recognize much shorter dsRNAs than 1 kbp.^24,40^

Interestingly, some studies claim that the presence of a 5’ end modification also affects MDA5’s dsRNA recognition in the same way as RIG-I.^12,25^ Based on previous studies and our observations, there seem to be three possible scenarios to explain why our prepared capped dsRNA avoids recognition by MDA5: (1) the presence of the cap structure contributes to the avoidance of dsRNA recognition not only by RIG-I but also by MDA5; (2) the basal level of MDA5 is too low to produce an effective response against dsRNA without an increase in its expression as happens during inflammation; or (3) it is simply too short; we prepared a duplex of 552 bp in length, and MDA5 only senses much longer substrates. Determining the exact answer to this question is important for fully understanding the exact mechanism of MDA5 activation; however, this is beyond the scope of the current study. Nevertheless, the observation that the cap structure almost completely reduced dsRNA immunogenicity is consistent with viral evasion strategies. Almost all RNA viruses have evolved the ability to add a cap structure to their RNAs.^28^ This takes advantage of cap-dependent translation, the main process of protein biosynthesis in mammalian cells, but also prevents immune recognition.

Importantly, knowing that we can disable RLR pathways by installing a 2’-*O*-methylated cap structure on dsRNA, we have an ideal tool to study the response of cell growth inhibitory pathways to dsRNA independent of cellular inflammation. Using this tool, we separated the actions of the OAS/RNase L and PKR pathways from that of the RLR-based cellular response. Both growth-inhibitory pathways responded effectively to dsRNAs even in the absence of cellular inflammation. OAS3 recognizes dsRNA and activates RNase L, leading to ssRNA degradation. Following dsRNA recognition, PKR phosphorylates eIF2α and represses global translation. Importantly, the degree of activation of both the pathways was the same in cells transfected with non-immunogenic and immunogenic dsRNAs. This means that even if dsRNAs can evade recognition by the RLRs during viral infection, the host cell is not powerless and is still able to take appropriate countermeasures to eliminate this threat. However, this observation raises the question: what is the benefit of activating RNase L and PKR without inducing cellular inflammation? Alternatively, is such a situation undesirable, and gives the cell no advantage in fighting potential threats? When analyzing our data, it is important to remember that dsRNAs are not only of viral origin but can also be endogenously derived. The higher abundance of dsRNAs in the cytoplasm of the cell may also be related to more frequent retrotransposition events, higher expression of long non-coding RNAs (lncRNAs), or the presence in the cytoplasm of more transcripts with long 3’UTRs that are prone to duplex formation.^1^ The question then arises as to whether it is necessary in such situations to induce inflammation and to warn the neighboring cells with cytokines and interferons that there is a serious threat to the homeostasis of the tissue or of the organism. Alternatively, it may be better to slow down cellular metabolism and take appropriate countermeasures to eliminate inappropriate levels of potentially dangerous transcripts; if this does not work, RNase L and PKR activity may eventually lead to cell apoptosis.^66–68^ If a mammalian cell chooses the latter scenario and tries to cope with dsRNA without involving other cells present in the tissue, the problem could be solved at the single-cell level without wasting the energy of the entire organism to deal with inflammation. Thus, the basal activity of OAS3 and PKR, which is maintained in cells growing under optimal conditions,^32^ appears to be an evolutionary adaptation to eliminate inappropriate levels of endogenously derived dsRNA, which poses no potential threat to neighboring cells in the tissue. Unfortunately, viruses have evolved a highly effective strategy to protect their duplexes from the primary cellular response of the host, which is particularly important for RNA viruses that replicate in the cytoplasm. Without a special shielding mechanism, viral dsRNA replication intermediates are fully exposed to host receptors, which prevents viruses from replicating efficiently. The strategy that probably all RNA viruses have developed is to restrict their replication to special factories called replication organelles (ROs).^69^ ROs are membrane organelles that isolate the site of viral replication from the host’s innate immunity, giving viruses a double protection, hiding replication intermediates that are not yet capped, as well as protecting the double-strandedness itself generated during replication.

Another strategy used by viruses to evade recognition of their transcripts by the host’s innate immunity is to introduce post-transcriptional modifications similar to those present in host RNA.^2,9,29^ Although the presence of these modifications in viral RNA was first characterized approximately half a century ago,^70,71^ the extent to which viruses benefit from their presence in their transcripts remains unknown. It is also possible that the presence of post-transcriptional modifications in viral RNA may be a side effect of viral replication in mammalian cells. Because all mammalian transcripts contain modified nucleotides, it is likely that viral transcripts would undergo the same post-transcriptional modifications as host RNA. Therefore, we tested how the three most common epitranscriptomic modifications in the human transcriptome, *i.e.* Ψ, m^6^A, and m^5^C, affect dsRNA-host cell interactions. Using *in vitro* transcription, dsRNAs carrying each of the three modified nucleotides were prepared. Unexpectedly, our data showed that the presence of any of the tested modifications did not significantly affect the dsRNA immunogenicity. This observation supports the notion that the 5’ end modification is a key player in the regulation of cellular inflammation in response to dsRNA. This appears to be the case at least in cells that rely on cytoplasmic dsRNA sensing. However, its effect on phagocytic cells, such as macrophages or dendritic cells (DCs), which recognize dsRNA not only through RLRs, but also through toll-like receptors (TLRs),^2,9,72^ has not been determined. Some clues come from our recent studies in which we transfected murine DCs (mDCs) with crude *in vitro* transcribed mRNA (IVT mRNA), *i.e.* mRNA containing double-stranded impurities generated by T7 RNA polymerase. We found that in mDCs, crude triphosphorylated IVT mRNA was more immunogenic than its cap1 version.^36^

Furthermore, using dsRNAs with a 2’-*O*-methylated cap structure and carrying post-transcriptional modifications, we examined the effects of Ψ, m^6^A, and m^5^C on cell growth inhibitory pathways. Despite previous suggestions that PKR activity may be regulated by post-transcriptional modifications,^47–49^ we observed no effect of any of these modifications on PKR-induced translational repression. As PKR forms higher-order assemblies in addition to dimers *in cellulo*,^56^ we also showed that none of the tested modifications affected PKR clustering on dsRNAs. Similarly, the immunogenic potential of dsRNA did not prevent PKR from forming higher-order assemblies. Interestingly, however, our immunofluorescence analysis showed that the presence of dsRNA foci in transfected mammalian cells is a side effect of lipofection, rather than a unique ability of dsRNA to form condensates inside living cells. Analogous foci were observed when ssRNA was delivered into A549 cells *via* lipofection.

Finally, we found that the presence of one of the post-transcriptional modifications tested, m^6^A, completely abolished dsRNA binding to OAS3 and thus did not induce RNase L-mediated ssRNA degradation. Nevertheless, we can extrapolate this observation to the other two OAS proteins, especially as it is OAS3 that has the highest affinity for dsRNA among the OAS proteins.^26,27^ Furthermore, after transfection of OAS3-KO cells with unmodified and m^6^A-modified dsRNA, only the former induced moderate rRNA degradation in this KO cell line, suggesting that OAS1/2 binding to dsRNA is indeed abolished by the presence of m^6^A. Interestingly, because dsRNAs containing m^6^A modification had no effect on translational repression in wild-type A549 cells, we agree that PKR is the key factor responsible for shutting down global host translation. However, it remains to be determined why this modification, among the three tested, has such an impact on OAS binding.

Using *in vitro* transcribed dsRNA, we have demonstrated that the key feature of dsRNA that regulates the onset of cellular inflammation in response to its recognition as a potential viral threat is its modification at the 5’ end. We showed that the presence of the cap0 structure alone significantly reduced the immunogenic potential of dsRNA. Furthermore, the introduction of dsRNA carrying a cap structure with only one 2’-*O*-methylation into mammalian cells did not induce any signs of inflammation. Nevertheless, mammalian cells can effectively respond to the potential threat of dsRNA, even without activating inflammatory pathways, through the action of OAS/RNase L- and PKR-dependent mechanisms. The answer to the question of why either of these two growth inhibitory pathways is fully functional in the absence of inflammation is that, in addition to coping with the viral threat, appropriate countermeasures should be taken when abnormal levels of endogenously derived dsRNA appear in the cytoplasm of the cell. In such situations, the expression of cytokines and interferons appears to be an unnecessary expenditure for the organism.

## Acknowledgments

We thank Susan Weiss for the A549 MAVS-KO, A549 OAS3-KO, A549 PKR-KO, and A549 RNase L-KO cell lines and Jacek Jemielity for the cap analogs. IFN-Beta_pGL3 was a gift from Nicolas Manel (Addgene plasmid # 102597). This study was supported by the National Science Centre (2021/42/E/NZ1/00314 to P.J.S.). The proteomic analysis was supported by an internal grant from the Medical University of Bialystok (Grant B.SUB.23.525). Next generation sequencing was performed thanks to Genomics Core Facility CeNT UW (RRID:SCR_022718), using NovaSeq 6000 platform financed by Polish Ministry of Science and Higher Education (decision no. 6817/IA/SP/2018 of 2018-04-10).

## Author contributions

K.D., J.C., and P.J.S. performed the majority of the experiments, with contributions from L.M.; A.P. analyzed the microscopy images under the supervision of A.B.; K.G. analyzed the RNA-Seq data; T.K. and D.C. conducted the protein mass spectrometry analysis; and P.J.S. conceived and directed the research and wrote the manuscript with contributions from all authors.

## MATERIALS AND METHODS

### Cell lines

A549 cell line from ATCC and A549-MAVS-KO, A549-OAS3-KO, A549-PKR-KO, A549-RNaseL-KO cell lines provided by Dr. Susan Weiss were maintained at 5% CO_2_ and 37°C in Dulbecco’s modified eagle’ medium (DMEM) supplemented with heat inactivated fetal bovine serum (FBS; 10% v/v), GlutaMAX (1% v/v) and penicillin/streptomycin (1% v/v).

### Plasmids

To generate plasmid encoding either sense or antisense strand for RNA duplex formation, part of *Gaussia* luciferase coding sequence was amplified *via* PCR using pJET_Gluc_A128 plasmid as a template, primers (Table S1), and Phusion™ High-Fidelity DNA Polymerase. The PCR amplicons were ligated into pJET1.2 plasmid utilizing CloneJET PCR Cloning Kit. Subsequently, pJET1.2 plasmids with cassette encoding sense or antisense RNA strand were digested with DraIII and XbaI and each insert was ligated into the DraIII/XbaI-treated pJET_Gluc_A128 with use of T4 DNA ligase, giving rise to the pJET_Gluc1 and pJET_Gluc2 constructs encoding sense and antisense strand of dsRNA, respectively.

### Antibodies

Rabbit anti-eIF2α, rabbit anti-phospho-eIF2α (Ser51), rabbit anti-GAPDH, rabbit anti-IFIT1, rabbit anti-MAVS, rabbit anti-OAS1, mouse anti-OAS2, rabbit anti-OAS3, rabbit anti-PKR, mouse anti-Puromycin, Rabbit anti-RNase L, and rabbit anti-Vinculin were used at 1:1000, whereas Goat anti-Mouse IgG (H+L) Secondary Antibody, HRP and Goat anti-Rabbit IgG (H+L) Secondary Antibody, HRP were used at 1:10000 forr western blotting. Mouse anti-G3BP1, rabbit anti-PKR, and mouse anti-Puromycin were used at 1:250, whereas Alexa Fluor® 488-AffiniPure Donkey Anti-Mouse IgG (H+L) and Donkey anti-Rabbit IgG (H+L) Highly Cross-Adsorbed Secondary Antibody, Alexa Fluor™ 488 were used at 1:1000 for immunofluorescence.

### *In vitro* transcription

RNA used in this study was obtained as described previously^36,38^ with some modifications. pJET-based construct linearized with PacQI restrictase served as a template for *in vitro* transcription. Reaction mixture contained RNA Pol buffer (40 mM Tris-HCl pH 7.9, 10 mM MgCl_2_, 1 mM DTT, 2 mM spermidine); 40 ng/μl of DNA template; mixture of nucleotides:

- CTP, GTP, UTP, 2 mM each, 0.5 mM ATP and 1.5 mM appropriate cap analog for capped RNA or 2 mM ATP (no cap analog) for non-capped triphosphorylated transcript to obtain unmodified transcripts;
- CTP, GTP, Pseudo-UTP, 2 mM each, 0.5 mM ATP and 1.5 mM appropriate cap analog for capped RNA or 2 mM ATP (no cap analog) for non-capped triphosphorylated transcript to obtain transcripts with Pseudo-UTP;
- 5-Methyl-CTP, GTP, UTP, 2 mM each, 0.5 mM ATP and 1.5 mM appropriate cap analog for capped RNA or 2 mM ATP (no cap analog) for non-capped triphosphorylated transcript to obtain transcripts with 5-Methyl-CTP;
- CTP, GTP, UTP, 2 mM each, 0.5 mM *N*6-Methyl-ATP and 1.5 mM appropriate cap analog for capped RNA or 0.5 mM *N*6-Methyl-ATP and 1.5 mM pppApG dinucleotide as transcription initiator for non-capped triphosphorylated transcript to obtain transcripts with *N*6-Methyl-ATP; 1 U/μl RiboLock RNase Inhibitor and home-made T7 RNA polymerase, 1 U/μl pyrophosphatase. Mixture was incubated at 37°C and after 2 h DNase I was added (0.1 U/μl) for additional 30 min. The crude RNA was purified using Monarch® RNA Cleanup Kit. Next, RNA was purified using Agilent 1260 Infinity HPLC system and XBridge Premier Oligonucleotide BEH C18 Column, 130Å, 2.5 µm, 4.6 mm X 150 mm HPLC was conducted at 55°C, with a linear gradient of buffer B (0.1 M triethylammonium acetate pH 7.0 and 50% acetonitrile) from 21.9% to 26.5% (for unmodified RNA and transcripts with Pseudo-UTP), 22.5% to 27.1% (for transcripts with 5-Methyl-CTP), or 24.0% to 28.6% (for transcripts with *N*6-Methyl-ATP) in buffer A (0.1 M triethylammonium acetate pH 7.0) over 30 min and a flow rate of 1 ml/min. RNAs from collected fraction was recovered by isopropanol precipitation. Non-capped RNAs from samples with capped RNAs were removed by two separate treatments, *i.e.* with RNA 5’ polyphosphatase and XRN-1, including purification using a Monarch® RNA Cleanup Kit step in between. Transcripts after enzymatic reactions were purified again utilizing Monarch® RNA Cleanup Kit.

### RNA labeling

Biotinylation or labelling with Cyanine5 was performed with poly(A) polymerase (PAP) as described previously.^60^ 10 μM HPLC-purified antisense RNA strand and 0.1 mM 2’-Azido-2’-dATP nucleotide analog were incubated in 20 μl with 1 U/μl RiboLock RNase Inhibitor, 2 μl of PAP enzyme, in 1x PAP buffer with 2.5 mM MnCl_2_, for 1 h at 37°C. RNA was purified using Monarch® RNA Cleanup Kit. “Click” reaction with 2 mM DBCO-PEG4-Biotin or 2 mM sulfo-Cyanine5 DBCO and RNA with incorporated 2’-Azido-2’-dATP analog was performed in 10% DMSO for 1 h at room temperature. Labeled RNA was purified using RNAClean XP magnetic beads. Labeling efficiency was verified either in agarose gel run in 1xTBE buffer and stained with EtBr or by RP-HPLC. RP-HPLC RNA samples were analyzed on Agilent 1260 Infinity HPLC system using an XBridge Oligonucleotide BEH C18 Column, 130Å, 2.5 µm, 4.6 mm X 50 mm, at 55°C, a linear gradient of buffer B (0.1 M triethylammonium acetate pH 7.0 and 50% acetonitrile) from 21.9% to 35% over 20 min and a flow rate of 0,9 ml/min.

### Double-stranded RNA preparation

Annealing of equimolar amounts of sense and antisense RNA strands was performed as follows: 360 ng/µl of each RNA strands was incubated in RNA annealing buffer (10 mM Tris–HCl pH 7.0, 150 mM NaCl, 1 mM EDTA) for 3 min in 90°C and then slowly cool down to room temperature. Efficiency of duplex formation was analyzed in agarose gel run in 1xTBE buffer and stained with EtBr. Additionally, formation of dsRNA was verified by RNase I assay as follows: 250 ng of single-stranded or double-stranded RNA was incubated in 10 µl in RNase I buffer (100 mM Tris–HCl pH 7.5, 10 mM NaCl, 0.1 mM EDTA) with 0.1 U of RNase I for 5 min at 37°C. Reaction products were analyzed in agarose gel run in 1x TBE buffer and stained with EtBr.

### RNA-Seq

A549 cells were seeded in 12-wells plate, cultured to 80% confluency, and transfected with 350 ng/ml of dsRNA using 2 µl mRNA Boost Reagent and 2 µl TransIT-mRNA Reagent (components of TransIT-mRNA Transfection Kit) in 100 µl Opti-MEM per 1 ml of cell culture. For mock transfection, cells were transfected respectively with no dsRNA. Four independent replicates were performed. Total RNA was isolated 5h post-transfection using TRI Reagent™ Solution and RNeasy Mini Kit. RNA libraries were prepared by the Genomics Core Facility (RRID: SCR_022718) at the University of Warsaw. Libraries were sequenced on Illumina NovaSeq 6000 with 2 × 150 bp paired-end reads. The raw sequence files were pre-processed using Trimmomatic v. 0.39 to trim Illumina adaptor sequences^73^ and low quality read fragments. Trimmed sequences were mapped to human reference genome provided by ENSEMBL v. grch38_snp_tran using Hisat2.^74^ Optical duplicates were removed using MarkDuplicates tool from GATK package v. 4.1.2.0.^75^ Mapped reads were associated with transcripts from Ensembl GRCh38 database v. 100 using HTSeq-count v. 0.9.1.^76^ Differentially expressed genes were selected using DESeq2 package v. 1.16.1.^77^ Fold change was corrected using normal method. Overrepresentation of Gene Ontology^78^ terms among the top 5% of genes (according to p-value) was assessed with clusterprofiler package.^79^

### dsRNA interactome capture

A549 cells were cultured to 90–95% confluency in 100 mm culture dishes and transfected with 350 ng/ml dsRNA as described above. Three independent replicates were performed. After 5 h cells were washed one-time in ice-cold PBS and 100 mm plate with cells covered with 2.5 ml of ice-cold PBS was deposited on a metal plate pre-cooled on ice and irradiated with 150 mJ/cm^2^ at 254 nm UV light in a UVP Crosslinker CL-1000. After crosslinking cells were detached with the use of a scraper in 600 μl of lysis buffer (50 mM Tris–HCl pH 7.4, 100 mM NaCl, 1% Igepal CA-630, 0.1% sodium dodecyl sulfate, 0.5% sodium deoxycholate supplemented with cOmplete™, EDTA-free Protease Inhibitor Cocktail) per dish. Cells and buffer from each dish were collected and transferred to one Eppendorf-type tube. The mixture was aspirated into a syringe and passed through a 26G needle seven times. Next, lysates were clarified by centrifugation 10 000 × g, at 4°C for 10 min. 10 µl of each lysate was saved for RT-qPCR analysis. Equilibrated in lysis buffer (buffer was exchanged twice) Dynabeads™ Protein G beads were coupled with mouse anti-dsRNA (J2) antibody (1 µg per 20 µl of beads) for 1 h at room temperature with rotation. Then beads were washed three times with lysis buffer. Lysates were incubated with 30 µl of antibody-coupled beads for 2 h at 4°C with rotation. Next, beads were washed with high salt wash buffer (50 mM Tris–HCl pH 7.4, 1 M NaCl, 1 mM EDTA, 1% Igepal CA-630, 0.1% sodium dodecyl sulfate, 0.5% sodium deoxycholate) twice, and twice with wash buffer (20 mM Tris–HCl pH 7.4, 10 mM MgCl_2_, 0.2% Tween-20). After last wash buffer was discarded, beads were frozen for MS/MS analysis.

Samples for MS/MS were prepared by adding 20 μl of 100 mM ammonium bicarbonate and 2.5 μl of 200 mM TCEP, vortexing, and then shaking at 10 000 rpm at room temaperature for 30 min. This was followed by adding 2 μl MMTS and shaking for additional 20 min at room temperature. A Trypsin/LysC Mix prepared in 8 M urea and 100 mM ammonium bicarbonate at a concentration of 0.02 μg/μl was added (50 μl per sample). Next, samples were incubated at 37°C with shaking for 4 hours, then supplemented with 300 μl ammonium bicarbonate and digested overnight. Acidification was done using 10 μl of 5% TFA. Peptides were purified using Oasis HLB 96-well plates, vacuum-dried, and resuspended in 60 μl of 2% acetonitrile and 0.1% TFA, then analyzed using an EvosepOne LC-MS setup coupled to an Orbitrap Exploris 480 mass spectrometer. For peptide loading, Evotips C18 trap columns were prepared according to the manufacturer’s protocol, including activation with 0.1% FA in acetonitrile, incubation in 1-propanol, and equilibration with 0.1% FA in water. Samples were loaded with 30 μl of 0.1% FA, and EvoTips were centrifuged at 600 × g for 1 min. Chromatographic separation was performed at 500 nl/min using a 44 min performance gradient on an EV1106 column. Data acquisition was in positive mode, with MS1 at 60,000 resolution and MS2 at 15,000 resolution, targeting the top 40 precursors with a dynamic exclusion of 20 s. Precursors were fragmented in HCD mode with a collision energy of 30%, and the system operated with a spray voltage of 2.1 kV and capillary temperature of 275°C

Raw data were analyzed with PEAKS Studio 10.6 64bit Bioinfor^80^ and searched against Uniprot human (78 120 entries) reference proteomes. Fixed modifications: methylthio (MMTS) at cysteines; variable: oxidation methionine, acetyl n-term. MS error and 0.1 Da, MS/MS level 0.2 Da, false discovery rate (FDR) 1%, digestion: trypsin semi-specific, max variable PTM per peptide: 3. Protein level analysis was performed using the ‘Label Free’ PEAKS module.

Each group consisted of three biological replicates; the average signal intensity was calculated for every condition. Data were analyzed in such a way as to indicate which proteins co-precipitate with the immobilized decoy in a repeatable and specific manner compared with the given control group.

The ratio of any protein identified and quantified was calculated in relation to the average level in the mock-treated samples normalized to ‘1.0’. Heat maps were done in GraphPad Prism. Gene ontology analysis was performed using https://geneontology.org.^81,82^

### RT-qPCR

A549 cells were seeded in 12-wells plate, cultured to 80% confluency, and transfected with dsRNA as described above. Three independent replicates were performed. After 5 h post transfection total RNA was isolated using TRI Reagent™ Solution and RNeasy Mini Kit. The residual DNA was removed by on-column treatment with DNase I. cDNA was synthesized using 1 µg of isolated RNA, M-MLV Reverse Transcriptase, and oligo-dT_20_ primer. 10 µl of cDNA was added to qPCR mixture containing SsoAdvanced Universal SYBR Green Supermix and gene-specific primers, listed in Table S1. Reactions were run in triplicate on LightCycler® 480 Instrument II (Roche).

Cell lysates (10 µl) from dsRNA interactome capture experiment were diluted with PBS and total RNA was isolated from them with NucleoSpin RNA Clean-up. Total RNA was subjected to DNase I treatment in dedicated buffer for 30 min in 37°C and purified again with NucleoSpin RNA Clean-up. cDNA synthesis (400 ng of RNA was used for reverse transcription) and qPCR analysis were performed as above.

### Immunoblotting

Cells were seeded in 24-well plates and cultured to 70% confluency, and transfected as above. After 24 h cells were lysed with RIPA buffer (50 mM Tris-HCl pH 8.0, 5 mM EDTA, 1% Igepal CA-630, 0.5% sodium deoxycholate, 0.1% sodium dodecyl sulfate, 150 mM NaCl) or with Luciferase Cell Culture Lysis Reagent, the later approach was used to study phosphorylation status of eIF2α. Lysates were mixed with Laemmli sample buffer and heated for 5 min at 95°C. Samples were separated by 10% SDS-polyacrylamide gel electrophoresis (PAGE). Then, proteins were transferred onto a nitrocellulose membrane using the Mini Trans-Blot® Cell and Criterion™ Blotter (Bio-Rad). Proteins on the membrane were stained with Ponceau S buffer, and 1 h blocking in 5% skim milk in PBST or in 5% bovine serum albumin in TBST was performed. Membranes were incubated with the appropriate primary antibody in PBST or TBST overnight at 4°C. After washing with PBST or TBST, membranes were incubated with secondary antibody in PBST or TBST for 1 h at room temperature. Detection was performed with the use of Pierce Fast Western Blot Kit in an Amersham Imager 600 (GE Healthcare Life Sciences)..

Puromycin incorporation assay was performed as described by Schmidt et al.^83^ Cells were seeded in 24-well plates and cultured to 70% confluency, and transfected as above. After 24 h, puromycin (4 µg/ml) was added to cells 30 minutes prior to harvesting in RIPA buffer. Lysates were mixed with Laemmli sample buffer and proceed as described above. After visualization proteins on membranes were stained with naphthol blue black solution (0.01% w/v).

### Pull-down

A549 cells were cultured to 90% confluency and incubated with 0 or 200 U/μl of IFNα for 5 h. For each condition (-/+IFNα) lysates from two 100 mm culture dishes were prepared. PBS-washed cells were detached with the use of a scraper in 800 μl of lysis buffer (20 mM Tris–HCl pH 7.4, 150 mM NaCl, 2 mM MgCl_2_, 2 mM DTT, 0.2% IGEPAL CA-630 supplemented with cOmplete™, EDTA-free Protease Inhibitor Cocktail) per dish. Cells and buffer were collected and transferred to one Eppendorf-type tube, and this mixture was aspirated into a syringe and passed through a 26G needle seven times. Next, lysates were centrifuged at 4°C, 10 000 × g, for 10 min. Lysates from both plates were pooled and then aliquoted. 100 µl of each lysate was saved as “input” for further analysis. Each of six aliquot was mixed with variously modified biotinylated dsRNA (no dsRNA added to control sample) and incubated at 4°C for 1 h with rotation. Then, pre-washed streptavidin magnetic beads were added, 10 μl of 50% slurry per sample, and incubated at 4°C for 30 min with rotation. Pre-washing of beads comprised three washes with lysis buffer (without protease inhibitors) followed by two washes with wash buffer (20 mM Tris–HCl pH 7.4, 150 mM NaCl, 2 mM MgCl_2_, 2 mM DTT, 0.2% Tween-20). After incubation with lysates, beads were washed twice with lysis buffer and twice with wash buffer. Beads and “input“ samples were mixed with Laemmli sample buffer and heated for 5 min at 95°C. Samples were separated in 10% SDS-PAGE. Then, proteins were transferred onto a nitrocellulose membrane using the Mini Trans-Blot Cell (Bio-Rad). Proteins on the membranes were stained with Ponceau S buffer, and 1 h blocking in 5% skim milk in PBST was performed. Membranes were incubated with the appropriate primary antibody in PBST overnight at 4°C. After washing with PBST, membranes were incubated with secondary antibody in PBST for 1 h at room temperature. Detection was performed as above.

### RNase L activity analysis

Cells were seeded in 12-well plates and cultured to 70% confluency, and transfected as above. Total RNA was isolated 24 h post transfection using TRI Reagent™ Solution and Monarch® RNA Cleanup Kit. Total RNA was analyzed in denaturing agarose gel (NBC buffer/formaldehyde) and stained with EtBr.

### Microscopy and image analysis

Cells were seeded on coverslips to reach 60% confluency the next day. Cells was transfected with Cy5-labelled dsRNA (350 ng per 1 ml of culture media) using 2 µl of mRNA Boost Reagent and 2 µl of TransIT-mRNA Reagent (components of TransIT-mRNA Transfection Kit) in 100 µl Opti-MEM per 1 ml of cell culture. After 5 h cells were fixed with 4% paraformaldehyde at RT for 10 min and permeabilized with 0.1% Triton-X100 in PBS at room temperature for 10 min. Next steps were: blocking for 1 h at room temperature with 5% donkey serum and 0.3 M glycine in PBS; staining overnight using primary antibody at 4°C followed by 3 washes with PBS; finally incubation with secondary antibody and 5 U of Alexa Fluor™ 555 Phalloidin conjugate for 1 h at room temperature. 1 µg/ml Hoechst 33342 in PBS was used to stain the nuclei for 5 min at room temperature, then cells were washed 2 times with PBS. Coverslips were mounted using ProLong™ Diamond Antifade Mountant and imaged using Carl Zeiss LSM 700 with Axio Imager Z2 confocal laser scanning microscope, with a 40x/1.3 oil objective. The Hoechst emission, Alexa 488 emission, Alexa 555 emission, and Cy5 emission were detected at emission spectra of 300-483 nm, 493-550 nm, 560-800 nm, and 644-800 nm, respectively, after excitation at 405 nm for Hoechst, 488 nm for Alexa 488, 555 nm for Alexa 555, and 639 nm for Cy5. Quantification of foci containing dsRNA or PKR was performed using Arivis Vision4D, Version 4.1.2. The analysis pipeline included following operations: denoising noise reduction by Gaussian filtering), intensity threshold, segmentation, size-based filtering, and spatial colocalization of segmented objects. Obtained results were next subjected to a statistical analysis using xxxxx test

### Luciferase assay

A549 cells were cultured to 80% confluency in 6-well plate and transfected with 1.5 µg IFN-Beta_pGL3 plasmid using Lipofectamine 3000. The following day transfected cells were seeded on 96-well plate. After 24 h cells were transfected with dsRNA (35 ng per 100 µl of culture media) using TransIT-mRNA Transfection Kit. 24 h post-dsRNA transfection cells were treated with Pierce™ Firefly Luc One-Step Glow Assay Kit and luminescence of firefly luciferase was measured in Synergy H1 (BioTek) microplate reader.

