## Supplemental Information for "Effective recognition of double-stranded RNA does not require activation of cellular inflammation"

**Table S1.** List of oligonucleotides used in this study.

| Name | Sequence | Purpose |
|---|---|---|
| GAPDH_for | | qPCR |
| GAPDH_rev | | qPCR |
| IFIT1_for | GATCAGCCATATTTCATTTTGAATC | qPCR |
| IFIT1_rev | GAAAATTCTCTTCAGCTTTTCTGTG | qPCR |
| IFIT2_for | AAGAGGAAGATTTCTGAAGAGTGC | qPCR |
| IFIT2_rev | TCTCCAAGGAATTCTTATTGTTCTC | qPCR |
| IFIT3_for | GAAGGAACTGGGCCGCCTGCTAAG | qPCR |
| IFIT3_rev | GCCCTGGCCCATTTCCTCACTACC | qPCR |
| IFNB1_for | GCCTGGACCATAGTCAGAGTG | qPCR |
| IFNB1_rev | AGCAATTGTCCAGTCCCAGAG | qPCR |
| MDA5_for | TTCCGCTATCTCATCTCGTGC | qPCR |
| MDA5_rev | GGCAGAAAGGTCAGGTAGTCC | qPCR |
| OASL_for | GTGCCTGAAACAGGACTGTTGC | qPCR |
| OASL_rev | CCTCTGCTCCACTGTCAAGTGG | qPCR |
| RIG-I_for | ATGTGCTCCTACAGGTTGTGG | qPCR |
| RIG-I_rev | ACACTGGGATCTGATTCGCAA | qPCR |
| Gluc1_for | GCCACGCTGTGTAATACGACTCACTATTAGGGAGTCAAAGTTCTGTTTG | Cloning of sense strand of dsRNA into pJET |
| Gluc1_rev | CGTCTAGACACCTGCACGTAGGGTCACCACCGGCCCCCTTG | Cloning of sense strand of dsRNA into pJET |
| Gluc2_for | CGTCTAGACACCTGCACGTAGGGAGTCAAAGTTCTGTTTG | Cloning of antisense strand of dsRNA into pJET |
| Gluc2_rev | GCCACGCTGTGTAATACGACTCACTATTAGGGTCACCACCGGCCCCCTTG | Cloning of antisense strand of dsRNA into pJET |

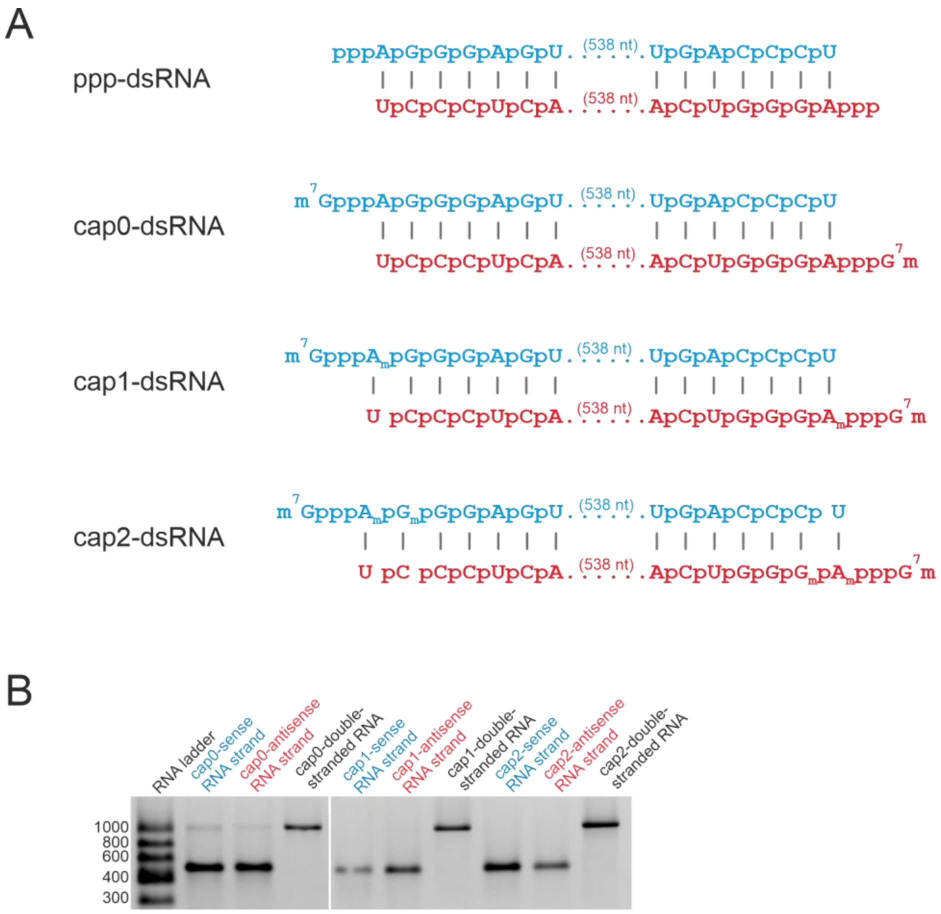


**Figure S1.** (A) Schematic representation of 5' end differently modified *in vitro* transcribed dsRNA. (B) Analysis of the sense and antisense dsRNA strands with cap0, cap1, or cap2 at the 5' end, and duplexes form with these strands on an agarose gel.

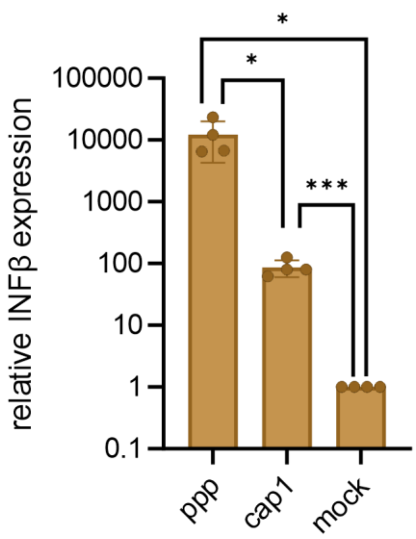


**Figure S2.** Levels of INFβ in A549 cells after 5 h transfection with ppp- and cap1-dsRNA. Bars represent the mean value of mRNA level change (relative gene expression) ± SD from four independent biological replicates, each independent biological replicate consists of a single transfection reaction. Each point represents data from one independent biological replicate. Data were normalized to mock treated cells. Statistical significance: * $P < 0.05$, *** $P < 0.001$ (one-way ANOVA with Turkey's multiple comparisons test). Only statistically significant differences were marked on the graph. Data were normalized to mock treated cells.

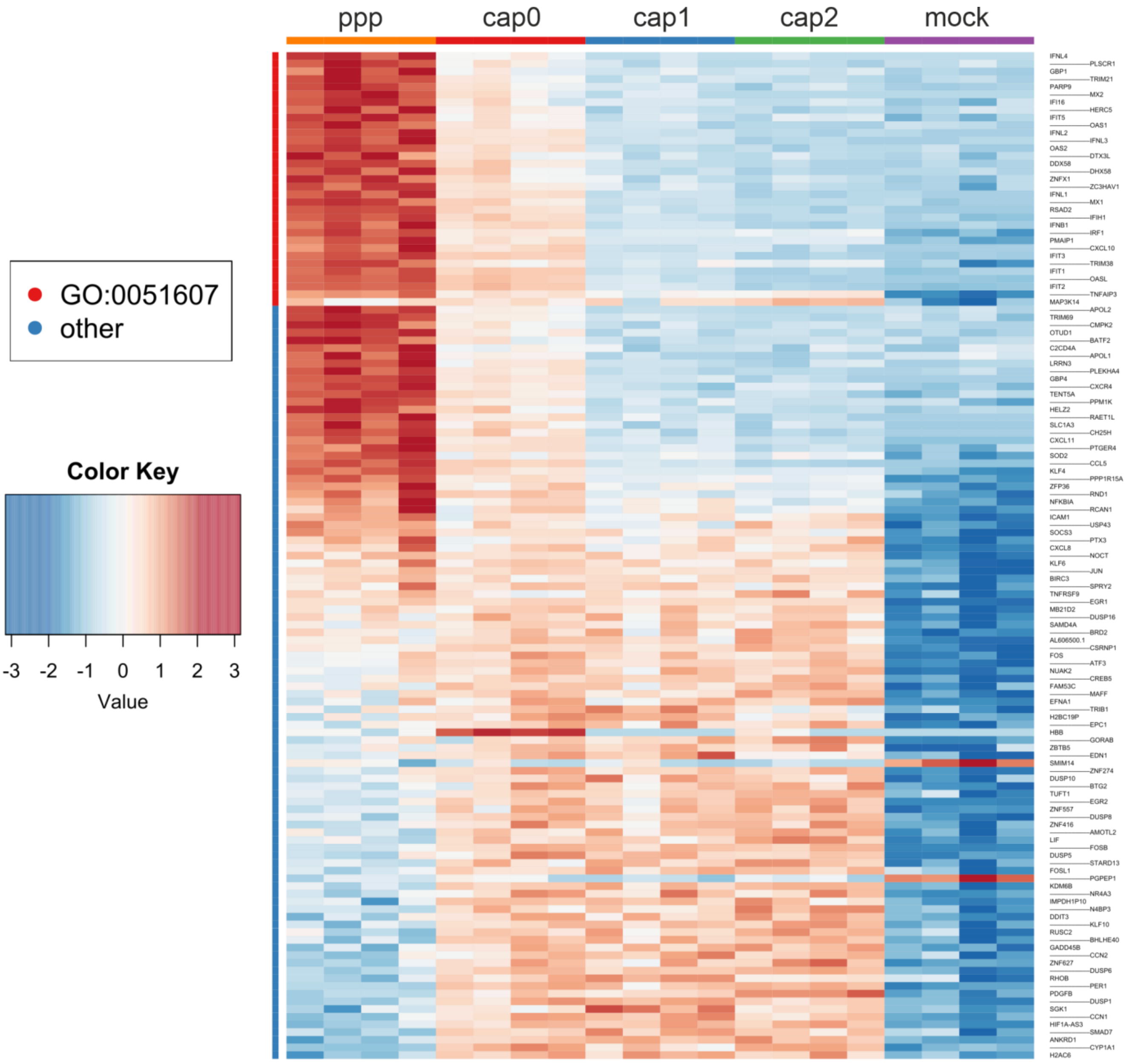


**Figure S3.** Heatmap of top 0.4% upregulated genes for all analyzed conditions (A549 cells were transfected for 5 h with dsRNA carrying different modifications at its 5' ends). Raw RNA-Seq data can be found in the supplemental files.

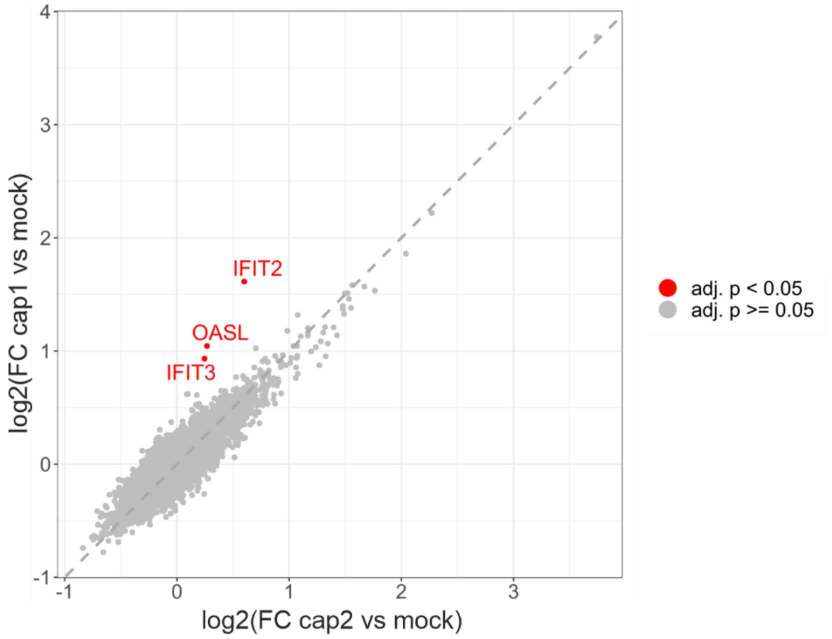


**Figure S4.** Scatter plot of log2-fold change (log2(FC)) upon dsRNA stimulation for 5 h. log2(FC) values in A549 cells transfected with cap2-dsRNA were plotted against those in A549 cells transfected with cap1-dsRNA. Three differentially expressed genes are colored red.

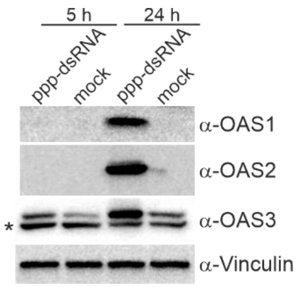


**Figure S5.** Comparison of ISG products level (Western blot analysis) in A549 cells after 5 h and 24 h transfection with ppp-dsRNA. * indicates unspecific band.

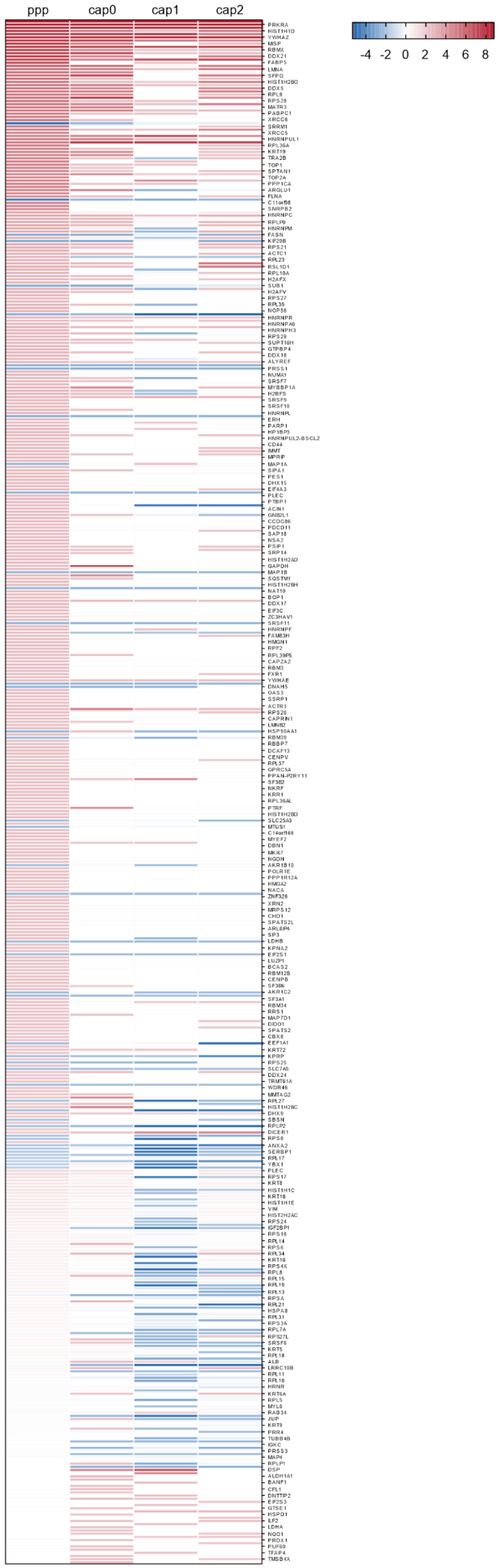
ppp
cap0
cap1
cap2
-4
-2
0
2
4
6
8
PRKRA
HIST1H1D
YWHAZ
MISP
RBMX
DDX21
FABP5
LMNA
SFPQ
HIST1H2BO
DDX5
RPL9
RPS28
MATR3
PABPC1
XRCC6
SRRM1
XRCC5
HNRNPUL1
RPL36A
KRT19
TRA2B
TOP1
SPTAN1
TOP2A
PPP1CA
ARGLU1
FLNA
C11orf98
SNRPB2
HNRNPC
RPLP0
HNRNPM
FASN
KIF20B
RPS21
ACTC1
RPL23
RSL1D1
RPL10A
H2AFX
SUB1
H2AFV
RPS27
RPL35
NOP56
HNRNPR
HNRNPA0
HNRNPH3
RPS29
SUPT16H
GTPBP4
DDX18
ALYREF
PRSS1
NUMA1
SRSF7
MYBBP1A
H2BFS
SRSF9
SRSF10
HNRNPL
ERH
PARP1
HP1BP3
HNRNPUL2-BSCL2
CD44
IMMT
MPRIP
MAP1A
SIPA1
PES1
DHX15
EIF4A3
PLEC
PTBP1
ACIN1
GNB2L1
CCDC86
PDCD11
SAP18
NSA2
PSIP1
SRP14
HIST1H2AD
GAPDH
MAP1B
SQSTM1
HIST1H2BH
NAT10
BOP1
DDX17
EIF3C
ZC3HAV1
SRSF11
HNRNPF
FAM83H
HMGN1
RPF2
RPL39P5
CAPZA2
RBM3
FXR1
YWHAE
DNAH5
OAS3
SSRP1
ACTR3
RPS26
CAPRIN1
LMNB2
HSP90AA1
RBM39
RBBP7
DCAF13
CENPV
RPL37
GPRC5A
PPAN-P2RY11
SF3B2
NKRF
KRR1
RPL36AL
PTRF
HIST1H2BD
SLC25A3
MTUS1
C14orf166
MYEF2
DBN1
MKI67
NGDN
AKR1B10
POLR1E
PPP1R12A
HMGA2
NACA
ZNF326
XRN2
MRPS12
CHD1
SPATS2L
ARL6IP4
SP3
LDHB
KPNA2
EIF2S1
LUZP1
BCAS2
RBM12B
CENPB
SF3B6
AKR1C2
SF3A1
RBM34
RRS1
MAP7D1
DIDO1
SPATS2
CBX8
EEF1A1
KRT72
KPRP
RPS25
SLC7A5
DDX24
TRMT61A
WDR46
MMTAG2
RPL27
HIST1H2BC
DHX9
SBSN
RPLP2
DICER1
RPS8
ANXA2
SERBP1
RPL17
YBX1
PLEC
RPS17
KRT8
HIST1H1C
KRT18
HIST1H1E
VIM
HIST2H2AC
RPS24
IGF2BP1
RPS18
RPL14
RPS6
RPL34
KRT16
RPS4X
RPL8
RPL15
RPL19
RPL13
RPSA
RPL21
HSPA8
RPL31
RPS3A
RPL7A
RPS27L
SRSF6
KRT5
RPL18
ALB
LRRC10B
RPL11
RPL10
HRNR
KRT6A
RPL5
MYL6
RAB34
JUP
KRT9
PRR4
TUBB4B
IGKC
PRSS3
MAP4
RPLP1
DSP
ALDH1A1
BANF1
CFL1
DNTTIP2
EIF2S3
GTSE1
HSPD1
ILF2
LDHA
NQO1
PRDX1
PUF60
TFAP4
TMSB4X

**Figure S6.** Protein specificity heatmap based on the data shown in Figure S7. This heatmap visualizes the specificity of identified proteins across different experimental conditions. The specificity value is calculated as the log2 intensity ratio compared to the control. All identified proteins with a specificity value greater than 0.0 in at least one of the conditions are shown.

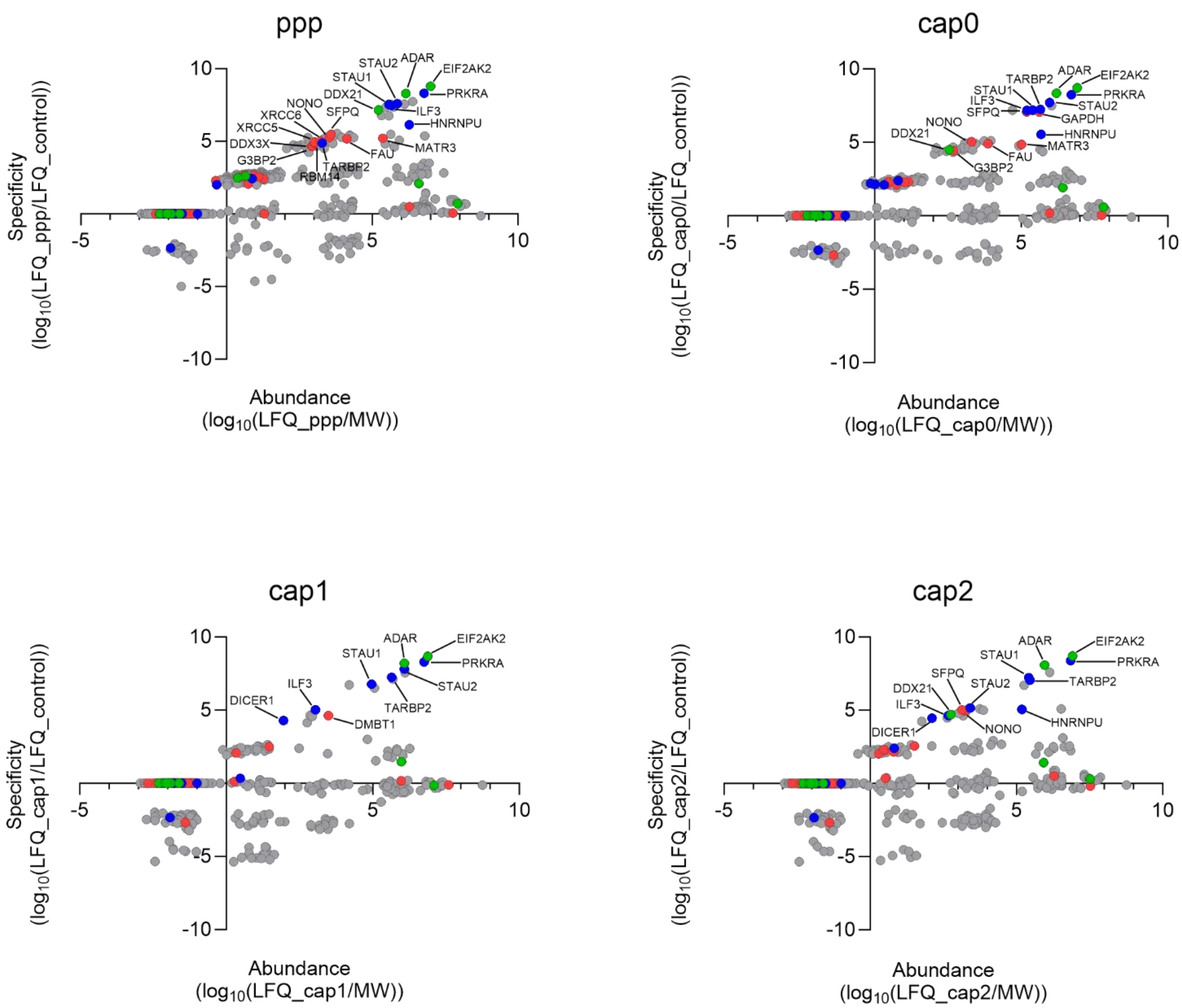


**Figure S7.** Protein abundance was quantified by normalized signal intensity (LFQ) relative to molecular weight. Specificity (enrichment) was determined by comparing LFQ intensities between dsRNA-transfected A549 cells and mock-treated cells. For proteins not detected in control samples, LFQ was arbitrarily set to 1 for calculations. Proteins annotated as "double-stranded RNA binding" (GO: 0003725) are shown in blue, proteins annotated as "innate immunity" (GO:0045087) are shown in red, and proteins belonging to both GO terms are shown in green.

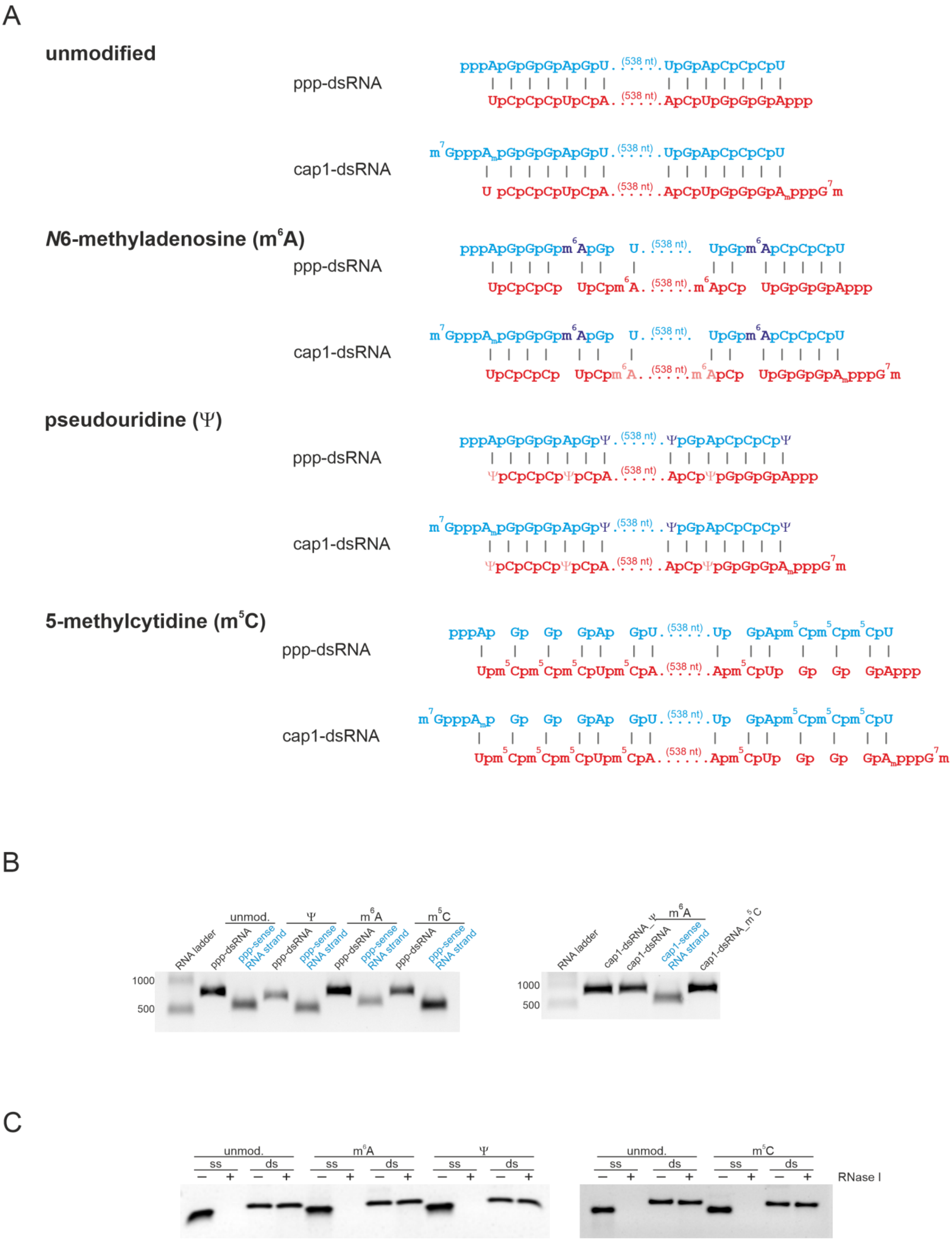


**Figure S8**. (A) Schematic representation of 5’ end differently modified *in vitro* transcribed dsRNA carrying $m^6A$, Ψ, or $m^5C$. (B) Analysis of the sense and antisense dsRNA strands with ppp at the 5’ end, and duplexes form with these strands on an agarose gel. (C) Stability analysis of the generated dsRNA with post-transcriptional modifications using RNase I. Transcripts were incubated with or without RNase I and the resulting products were analyzed on an agarose gel.

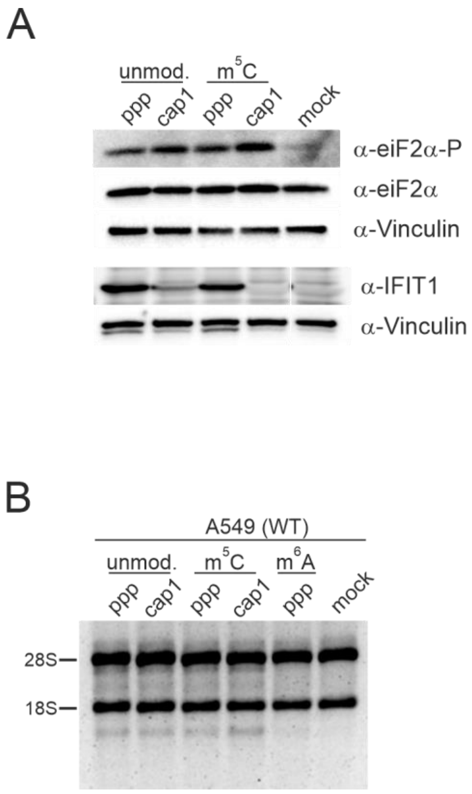


**Figure S9.** (A) Activation of RLR and PKR pathways is not affected by the presence of $m^5C$ within dsRNA. A549 cells were transfected with ppp- and cap1- dsRNA or with its counterparts carrying $m^5C$ for 24 h and IFIT1 expression level as well as phosphorylation status of eIF2α were assessed using Western blot. (B) The presence of $m^5C$ within dsRNA does not affect RNase L activity. RNase L activity in A549 cells assessed by rRNA integrity. Total RNA was isolated after 24 h transfection with post-transcriptionally modified ppp- or cap1- dsRNA, and analyzed on a 1 x TBE agarose gel.

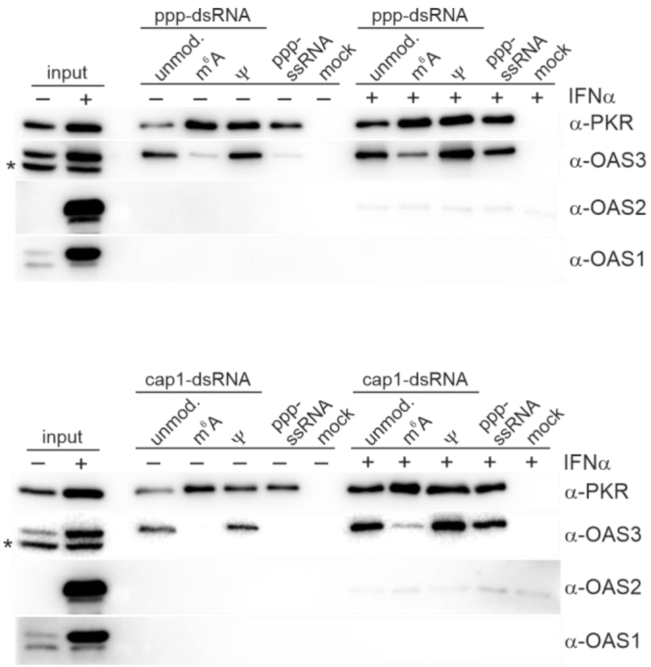


**Figure S10.** Co-purification of endogenous proteins from lysates of IFNα-treated (200 U/ml) and untreated A549 cells with biotinylated ppp- or cap1-dsRNA. PKR, OAS3, OAS2, and OAS1 were detected in precipitates by wester blotting. * indicates unspecific band.

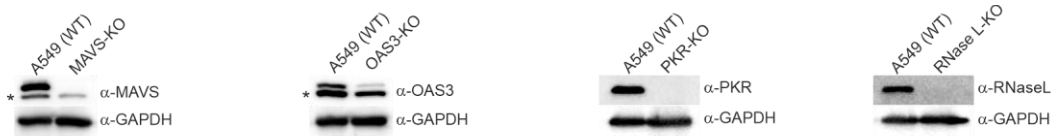


**Figure S11.** Verification of MAVS, OAS3, PKR and RNase L knocked out in MAVS-KO, OAS3-KO, PKR-KO and RNase L-KO cells, respectively, using western blotting. * indicates unspecific band.

A

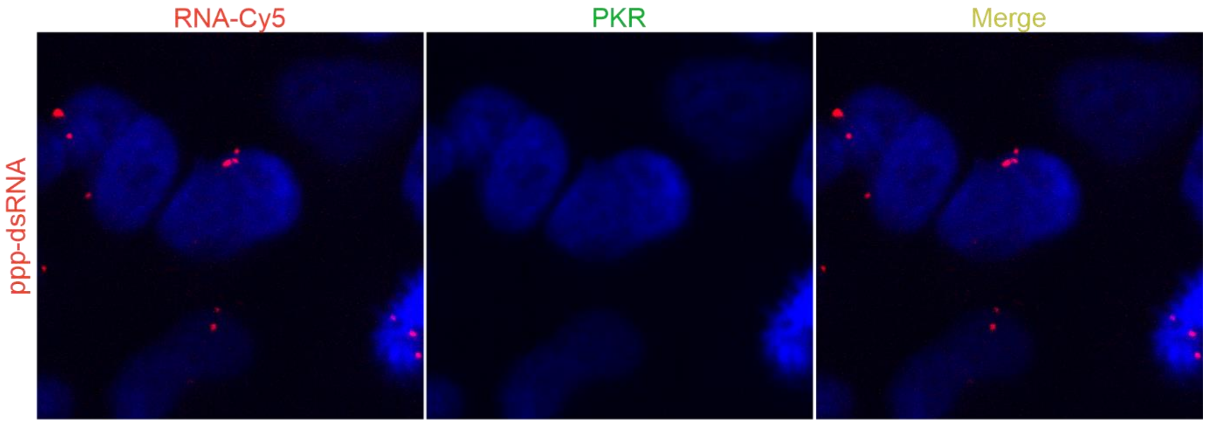


B

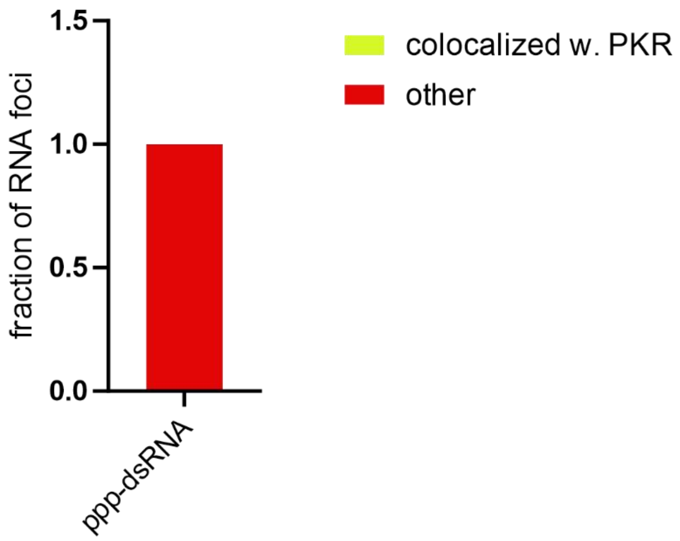


**Figure S12.** PKR is dispensable for RNA foci formation. (A) Immunofluorescence analysis of PKR and Cy5-labeled ppp-dsRNA in PKR-KO cells. (B) Quantification of Cy5-labeled ppp-dsRNA colocalization with PKR (34 cells were analyzed). Presented bar represents all foci counted, yellow color represents the fraction of foci in which RNA colocalized with PKR, whereas red color represent the fraction of foci in which only signal from PKR was observed.
